# Identification of a novel HASPIN inhibitor and its synergism with the PLK1 inhibitor

**DOI:** 10.1101/2022.09.01.506282

**Authors:** Eun-Ji Kwon, Karishma K. Mashelkar, Juhee Seo, Yoonze Shin, Kisu Sung, Sung Chul Jang, Sang Won Cheon, Haeseung Lee, Byung Woo Han, Sang Kook Lee, Lak Shin Jeong, Hyuk-Jin Cha

## Abstract

**Background:** HASPIN, a mitotic kinase for Histone H3, is a promising target for anti-cancer therapy. However, as HASPIN is an atypical kinase with low similarity to eukaryotic protein kinases, development of a HASPIN inhibitor from the conventional pharmacophore of kinase inhibitors would be technically challenging.

**Methods:** Chemical modifications of a cytotoxic 4’-thioadenosine analogue with high genotoxicity and multiple kinomescan profiles were performed to produce a novel non-genotoxic kinase inhibitor, LJ4827. The mode of action of this inhibitor with clear anti-cancer activity was inferred based on transcriptomic and chemical similarity to known drugs.

**Results:** The specificity and potency of LJ4827 as a HASPIN inhibitor were validated by *in vitro* kinase screening and subsequent X-ray crystallography. As predicted, LJ4827 treatment delayed mitosis by clear inhibition of the recruitment of Aurora B at the centromere in cancer cells, without a genotoxic response. Through transcriptome analysis of lung cancer patients, *PLK1* was predicted as a druggable synergistic partner to complement HASPIN inhibition. Cotreatment with the PLK1 inhibitor BI2536 and LJ4827 led to pronounced cytotoxicity of lung cancers *in vitro* and *in vivo*.

**Conclusion:** Simultaneous inhibition of both HASPIN and PLK1 is a promising therapeutic strategy for lung cancers.

## Introduction

As there is a strong correlation between the active cell cycle and the degree of malignancy, pathologists use the mitotic index of cancer specimens for diagnosis of cancer [1]. While many conventional chemotherapeutics directly or indirectly impede the cell cycle of cancer cells by triggering cell cycle checkpoints [2, 3], the side effects of these chemotherapeutics on the normal cell cycle in actively renewing tissues (e.g., bone marrow, hair follicles, and gastrointestinal tract) have repeatedly resulted in discouraging clinical results. Thus, for identification of druggable targets to exclusively perturb the cancer cell cycle or as a synthetic lethal partner of oncogenic mutations, the cancer cell cycle has been extensively characterized [4–6].

HASPIN (Haploid Germ Cell□Specific Nuclear Protein Kinase), encoded by germ cell-specific gene 2 protein (*GSG2*), directly phosphorylates the threonine 3 residue of Histone H3 (H3T3ph) from late G2 throughout anaphase in mitosis [7]. The H3T3ph then serves as a docking site for centromere localization of the chromosome passenger complex (CPC), which tightly specifically controls proper kinetochore microtubule attachment mediated by Aurora B kinase [8, 9]. Thus, depletion of HASPIN causes chromosome misalignment, premature chromatid separation, and mitotic delay in somatic cancer cell models [7, 8]. While other key mitotic kinases, of which depletion causes severe phenotypic and developmental abnormality [e.g., CDK1 [10], PLK1 [11] or Aurora A [12]], lack of HASPIN in the mouse reveals normal physiology (except for testicular abnormality) [13], and knockout of HASPIN in embryonic stem cells (ESCs) maintains normal mitosis [14]. These studies imply HASPIN as a promising mitotic target to exclusively impede cancer mitosis [15].

The majority of kinase inhibitors compete with ATP by binding in or around the ATP binding cleft and are classified as type I, type II, type III, or type IV depending on the binding mode [16]. In particular, the conserved ATP/Mg^2+^ binding motif, Asp (D)-Phe (F)-Gly (G) residues of the DFG motif, is responsible for reversible confirmation of active or inactive kinases. Thus, type I and II inhibitors, comprising most of the currently known kinase inhibitors, have been designed to lock the confirmation of the ATP pocket in ‘DFG-in’ (type I) or ‘DFG-out’ (type II), respectively [16]. Notably, HASPIN is classified as an atypical eukaryotic kinase (ePK), with Asp-Tyr-Thr (DYT) instead of the DFG motif [17] and shares low sequence homology with other ePKs [17]. A few HASPIN inhibitors have been reported with discrete chemical scaffolds (e.g., imidazopyridazine CHR-6494, nucleoside 5ITU, the β-carboline acridine LDN-211898) [18].

Identification of the mode of action (MoA) of a new compound is an essential but challenging step in the drug discovery process [19]. Recently, advanced computational approaches based on large data of the drug transcriptome [20], chemical structures [12], and text mining [21], have been used to elucidate MoA. In particular, based on a large dataset of the transcriptome of compounds (with known MoA) [20], a new compound of interest can be connected to the gene signatures of a subset of known drugs by matching of compound-specific expression profile, which leads to prediction of similarities in MoA [22].

Herein, among three chemical derivatives modified from the cytotoxic 4’-thioadenosine analogue LJ4425, a cytotoxic multi-kinase inhibitor [23], we identified the novel potent HASPIN inhibitor LJ4827 through computational analysis of the drug-induced transcriptome and subsequent kinomescan profiling. Mitotic delay and inhibition of CPC action were evident without DNA damage, unlike other HASPIN inhibitors. More importantly, PLK1 was further determined as a synergistic partner of HASPIN inhibition based on patient tumor transcriptome and survival data. The anti-cancer potential of simultaneous inhibition of HASPIN and PLK1 was validated by *in vitro* and *in vivo* assays of lung cancer, leading to the proposal of a novel anti-cancer therapeutic strategy.

## Materials and Methods

### Generation and preprocessing of RNA sequencing data

Total RNA was isolated from HeLa cells before and after treatment with vehicle, LJ 4827, or 5ITU in duplicates using Trizol according to the manufacturer’s instructions. For library construction, we used the TruSeq Stranded mRNA Library Prep Kit (Illumina, San Diego, CA). Briefly, the strand-specific protocol included the following steps: (1) strand cDNA synthesis, (2) strand synthesis using dUTPs instead of dTTPs, (3) end repair, A-tailing, and adaptor ligation, and (4) PCR amplification. Each library was then diluted to 8 pM for 76 cycles of paired-read sequencing (2 × 75 bp) on an Illumina NextSeq 500 following the manufacturer’s recommended protocol. Read quality was assessed using FastQC (v) and poor-quality bases (Phred score < 20) were eliminated using TrimGalore (v0.6.6). Trimmed reads were aligned to the human reference genome (GRCh38) using the STAR aligner (v2.7.9a) with default parameters. Gene-level expression values such as transcripts per million (TPM) and read counts were calculated using RSEM (v1.3.3.) with human gene annotation (GRCh38.84). FASTQ format files, gene-level count data, and TPM of all samples are available in the Gene Expression Omnibus. Among the xxx genes, xxx protein-coding genes were utilized for subsequent analysis. Differential gene expression analysis was performed using the ‘DESeq2’ package (v3.15) in R (v4.2.1).

### CMap analysis to infer MoA

CMap provides a web-based tool for searching for compounds that give rise to similar or dissimilar expression signatures to an input signature, referred to as a set of differentially expressed genes. It contains an extensive catalog of transcriptome profiles for 9 core human cell lines (A375, A549, HA1E, HCC515, HEPG2, HT29, MCF7, PC3 and VCAP) before and after treatment with each of 29,679 small molecules. To infer the MoA of LJ 4827, we queried CMap reference database for differentially expressed genes induced by LJ 4827 treatment (DEGs, top or bottom 150 genes selected based on Wald statistic values of DESeq2 results) and obtained a list of compounds that induce similar expression signatures to LJ 4827 (normalized connectivity score > 2).

### Chemical similarity of compounds

Simplified Molecular Input Line Entry System (SMILES) of 18 top-scoring compounds and LJ 4827 were obtained from ChEMBL (https://www.ebi.ac.uk/chembl/). Structural similarity between compounds was determined by Tanimoto coefficients calculated using RDKit (v2022.03.1) with default settings in Python3 (v3.9).

### Functional enrichment analysis

Quantitative changes in gene expression levels between groups (treatment vs vehicle, tumor vs normal) were estimated by using the ‘DESeq2’ package in R. Differentially expressed genes were selected with cutoffs of false discovery rate (FDR) adjusted P-value < 0.01 and |fold change| ≥ 2. Over-representation analysis of Gene Ontology (GO) among DEGs of LJ4827 and 5ITU was performed using the R package topGO. Gene set enrichment analysis (GSEA) was performed with the REACTOME database based on a list of genes ranked by Wald statistic from DESeq2 result via the ‘msigdbr’ (v7.5.1) and ‘fgsea’ (v1.22.0) package in R.

### The Cancer Genome Atlas (TCGA) transcriptome data analysis

Transcriptome (TPM) and clinical information of 9563 cancer patients in Pan-Cancer Atlas were obtained from the UCSC Xena Functional Genomics Explorer (https://xenabrowser.net/) and cBioPortal (https://www.cbioportal.org/), respectively. Active mitosis signature enrichment score (AMSES) of each patient was computed by single-sample GSEA (ssGSEA) analysis using the active mitosis signature. The ssGSEA was conducted through R packages ‘fgsea’ based on patient-wise z-transformed TPM values of all genes. Survival analysis was conducted to test the difference in the survival rate in patients between two groups, high and low, determined based on GSG2 expression level or Active mitosis signatures score. The hazard ratio (HR) and P-value were estimated from Cox proportional hazards regression analysis and the log-rank test by using the package ‘survival’ (v3.4-0) in R. The synthetic lethal (SL) partner of CSG2 was explored within the active mitosis signature by using tumor transcriptome data of TCGA LUAD patients (N=508). Among the signature, genes encoding kinases were considered candidates for SL partners due to their druggability. Patients were divided into GSG2-high and GSG2-low groups based on *GSG2* expression. Association between kinase gene expression and overall survival rates in the GSG2-low group or GSG2-high group were estimated using a Cox regression analysis.

### Immunoblotting and immunofluorescence

Cells were lysed with tissue lysis buffer supplemented with 0.2 mM sodium vanadate and 1 mM protease inhibitor cocktail (Roche, Basel, Switzerland), and for immunoblotting. For immunofluorescence, cell were fixed with 4% PFA for 10 min at RT followed by permeabilization with 0.1% Triton X-100 for 2 min and blocking with 3% BSA for 1 hr at RT. Antibodies for Cyclin B1 (#sc-245) and β-actin (#sc-47778), were purchased from Santa Cruz Biotechnology. Antibodies for pH2AX (#9718S), pH3ser10 (#9710S), CENP-F(#58982S), and cleaved caspase-3 (#9664S) were purchased from Cell Signaling Biotechnology. Antibodies for pH3T3(ab222775) and AURKB (ab3609) were purchased from Abcam. HASPIN (NBP1-26626) was purchased from Novus Biologicals.

### Cloning, expression, and purification of HASPIN for structure determination

The kinase domain of human HASPIN (residue 452 – 798) was cloned into the expression vector pET-21a(+) (Novagen, Madison, WI, USA). The genomic DNA of HASPIN was provided from the Korea Human Gene Bank, Medical Genomics Research center, KRIBB, Korea. The plasmid that contains the kinase domain of human HASPIN was transformed into *Escherichia coli* strain Rosetta™ 2(DE3) pLysS (Novagen, Madison, WI, USA). The transformed cells were grown at 37□ until OD_600_ reached 0.8, and induced with 0.5 mM isopropyl β-D-1-thiogalactopyranoside (IPTG). After incubation for additional 18 h at 20□, the cells were harvested and resuspended using lysis buffer [20 mM Tris-HCl, pH 7.5, 500 mM NaCl, 35 mM imidazole, and 1 mM phenylmethanesulfonylfluoride]. For the affinity chromatography, 5 mL HiTrap™ chelating HP column (GE Healthcare, Chicago, IL, USA) charged with Ni^2+^ was used. The column was equilibrated with buffer A [20 mM Tris-HCl, pH 7.5, 500 mM NaCl, and 35 mM imidazole] and eluted with buffer B [20 mM Tris-HCl, pH 7.5, 500 mM NaCl, and 1 M imidazole]. The eluent was loaded into HiLoad 16/600 Superdex75™ pg column (GE Healthcare, Chicago, IL, USA) that was pre-equilibrated with buffer C [20 mM Tris, pH 7.5, 250 mM NaCl, 10% glycerol, and 1 mM 1,4-dithiothreitol]. The purified HASPIN protein was finally concentrated to 5 mg/mL for crystallization experiments.

### Crystallization, data collection, and structure determination of HASPIN in complex with LJ4827 and with LJ4760

LJ4827 and LJ4760 were respectively added to the purified HASPIN protein with molar ratio of 1:3 and were co-crystallized by the sitting-drop vapor diffusion method at 22□. Diffraction-quality crystals were obtained at the reservoir containing 23.75% PEG 4000, 0.2 M ammonium sulfate, and 0.1 M sodium acetate at pH 4.6 for LJ4827, and 23.75% PEG4000, 0.325 M ammonium sulfate, and 0.1 M sodium acetate at pH 4.6 for LJ4760. Crystals were cryoprotected in the crystallization solution supplemented with 30% glycerol and flash-frozen into liquid nitrogen. X-ray diffraction data were collected at the Eiger 9M detector (Dectris Ltd., Baden, Switzerland) at the beamline 5C of the Pohang Light Source, Korea. HKL2000 program suite was used for raw data indexing and scaling [24]. The structures of the kinase domain of HASPIN in complex with LJ4827 and with LJ4760 were solved by the molecular replacement method using the HASPIN structure with PDB ID 2WB8 [25] as a reference model in the PHENIX Phaser-MR program [26]. The structures were further refined by Coot in the CCP4i program suite [27] and phenix.refine in PHENIX [26].

### Docking studies

AutoDock Vina [28] was used for docking studies. The HASPIN structure in complex with LJ4827 was used for the control docking experiment and the grid box sized with 10×10×10 points was centered at the LJ4827 binding site with 1.0 Å spacing. LJ4760, 5ITU, CHR6494 were used as ligands for docking and the structures of ligands were obtained using PRODRG [29].

### *In Vivo* Tumor Xenograft Model

All animal experiments were conducted according to the guidelines approved by the Seoul National University Institutional Animal Care and Use Committee (IACUC permission number SNU-220217-1). Balb/c-nu mice (male, 4-weeks-old; OrientBio, Seoul, Korea) were allowed one-week acclimation prior to the experiment. A549 cells (4.5 × 10^6^ cells in 200 μL PBS) were injected subcutaneously into the flanks of mice, and tumors were allowed to grow for 14 days until their volume reached approximately 70 mm^3^. The mice were randomly divided into four groups for vehicle control and treatment groups (n = 5); vehicle control (DMSO: cremophor: normal saline = 1:1:18), LJ4827 (1 mg/kg body weight), BI2536 (10 mg/kg body weight), or a combination of LJ4827 (1 mg/kg body weight) and BI2536(10 mg/kg body weight). LJ4827 compound was dissolved in vehicle (DMSO: cremophor: normal saline = 1:1:18) and BI2536 compound was dissolved in vehicle (polyethyleneglycol-400: normal saline = 1:4). LJ4827 and vehicle control (DMSO: cremophor: normal saline = 1:1:18) were intraperitoneally administered three times per week. BI2536 was intraperitoneally administered two times per week, and a combination of LJ4827 and BI2536 were administered each three times per week and two times per week for 23 days. The body weight and tumor volume were measured every 2-3 days. The tumor volume was measured using a digital slide caliper according to the following formula: tumor volume (mm^3^) = 0.52 × (width□×□length ×□height).

### Statistical analysis

The quantitative data are expressed as the mean values ± standard deviation (SD). Student’s unpaired t-tests for two groups or one-way ANOVA following Tukey mulpile comparison, was performed to analyze the statistical significance of each response variable using the PRISM. p values less than 0.05 were considered statistically significant (*, p < 0.05, **, p < 0.01, ***, p < 0.001, ****, p < 0.0001 and n.s for not significant).

## Results

### Chemical modification of genotoxic 4’-thioadenosine analogs

Previously, we reported a cytotoxic multi-kinase inhibitor LJ4425 with the 4’-thioadenosine structure [23] (Fig. 1A). However, further development of LJ4425 as an anti-cancer drug was discontinued due to high toxicity in animal models (data not shown). We speculated that the high toxicity of LJ4425 resulted from genotoxicity induced by interfering with DNA elongation due to the existence of a 5’-hydroxyl group, similar to other anti-cancer nucleoside analogues such as gemcitabine and clofarabine (Fig. 1B). To rule out this possibility, three novel compounds that replaced the 5’-hydroxyl group with amine (LJ4760), azide (LJ4827), and urea (LJ4857) moieties (Fig. 1C) were synthesized (Scheme S1). Briefly, D-ribose was converted to the key intermediate **1** by a known method [21]. Prior to modification of the 5′-hydroxyl group, the adenine moiety of **1** was protected with *N,N*-di-Boc, and the 5′-hydroxyl derivative **2** was produced after removal of the TBDPS group. Mesylation of **2** followed by treatment with sodium azide afforded the 5′-azido derivative **3**, which was hydrolyzed with 50% aq. formic acid to give the azido derivative **4** (LJ4827). Reduction of **4** with PPh_3_ and H_2_O afforded the amino derivative **5** (LJ4760). For synthesis of the urea derivative **8** (LJ4857), the 5′-azido group of **3** was reduced to the 5′-amino group to produce **6**, which was successively treated with trichloroacetyl isocyanate and methanolic ammonia to afford the 5′-urea derivative **7**. Removal of the acetonide-protecting group of **7** under acidic conditions produced the urea derivative **8** (LJ4857). As an initial prediction, DNA damage response, determined by phosphorylated H2AX (serine 139), a marker for double strand break (DSB), of A549 cancer cells after treatment with LJ4827 or LJ4760 (but not LJ4457) was significantly weakened compared to that of LJ4425 (Fig. 1D). Despite the lack of genotoxicity, LJ4827 was likely to efficiently inhibit cell growth of multiple cancer cell lines (Fig. 1E).

**Figure. 1.**
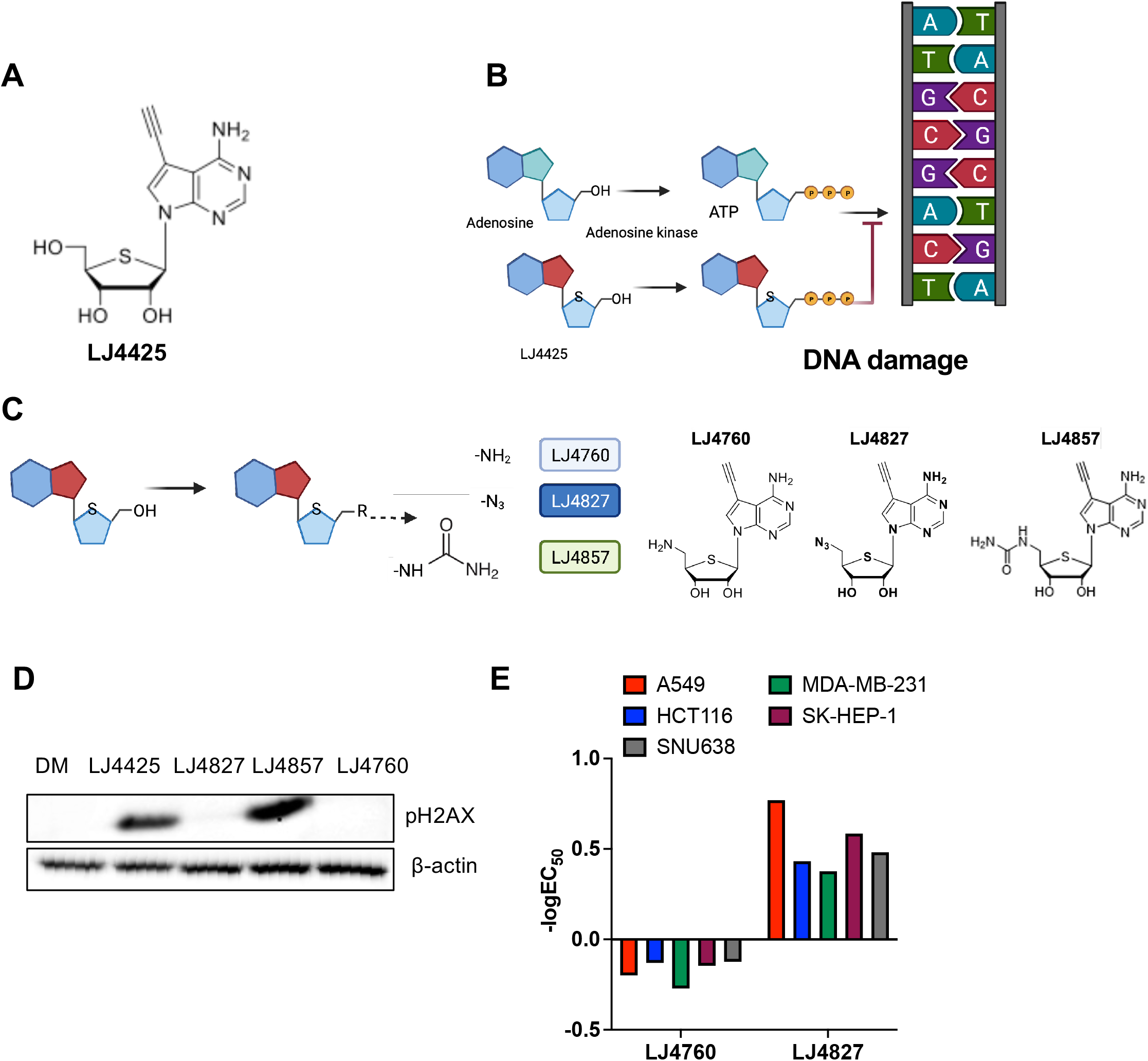
Chemical modification of genotoxic 4’-thioadenosine analogs. (A) Chemical structure of LJ4425, (B) Graphical presentation of Adenosine kinase analogue, LJ4425, interfering in DNA elongation due to the existence of 5’-hydroxyl group (C) Derivatives of LJ4425 by replacing 5’-hydroxyl group to amine (LJ4760, left), azide (LJ4827, middle) and urea (LJ4857, right) (E) Immunoblotting analysis for pH2AX on HeLa cells after each derivatives treatment (500nM), β-actin for equal protein loading control (F) Graphical presentation of EC_50_ value of LJ4760 and LJ4827 on cell growth inhibition of six different cell lines

### Transcriptome-guided MoA inference of LJ4827

The potency of LJ4827 for cell growth inhibition of multiple cancer cell lines (Fig. 1E) urged further examination of the MoA. To this aim, we profiled LJ4827-induced differentially expressed genes (DEGs) via RNA sequencing and queried DEGs against the CMap database to search for compounds that resulted in similar expression changes [22] (Fig. 2A). Among the top-scoring compounds, adenosine kinase (AdK) inhibitors followed by PKA, IKK, CDK, and JNK inhibitors were highly enriched (Fig. 2B). Based on the assumption that chemically similar drugs are likely to share common targets or MoA [30], chemical structural similarity between LJ4827 and the top-scoring compounds was determined by Tanimoto coefficient (Fig. 2C). The AdK inhibitor 5-iodotubercidin (5ITU) was most similar to LJ4827 with respect to perturbing effects on the transcriptome and chemical structure (Fig. 2D). In order to explain the anti-cancer effect of LJ4827 in multiple cancers (Fig. 1E) with AdK inhibition by LJ4827, the catalytic activity of AdK was assessed *in vitro* with 100 nM or 500 nM of LJ4827, where a clear anti-cancer effect was observed (data not shown). However, unexpectedly, AdK inhibition by either 5ITU or LJ4827 was marginal compared to the positive control AdK inhibitor (Fig. 2E), suggesting that the anti-cancer effect of LJ4827 is independent of AdK inhibition.

**Figure. 2.**
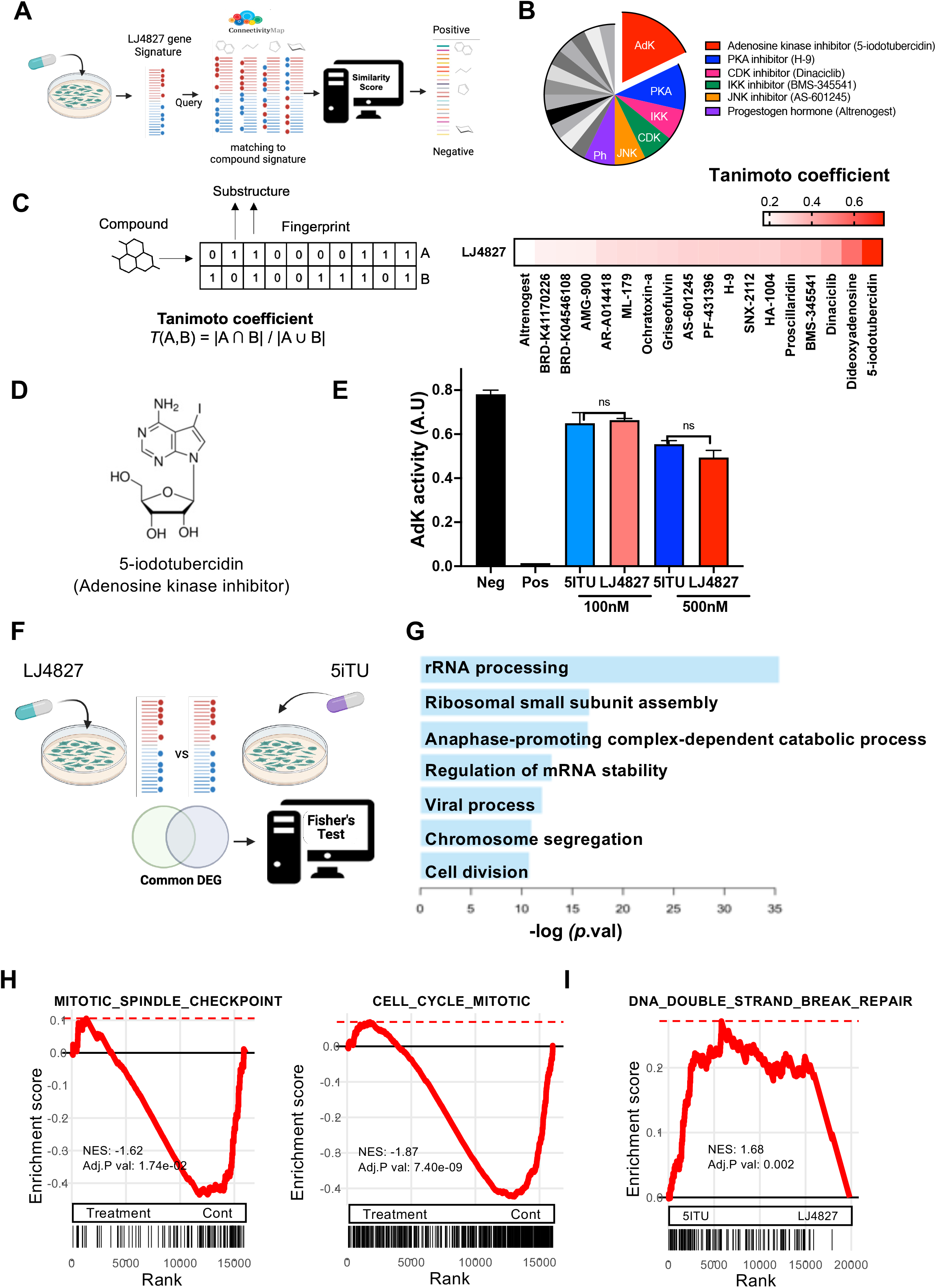
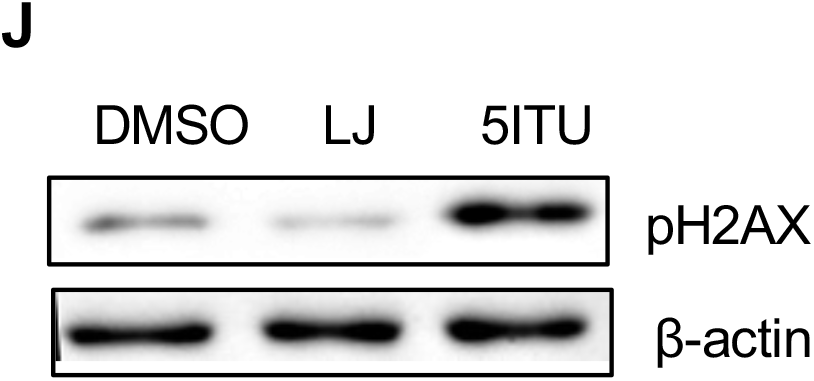
Transcriptome-guided MoA inference of LJ4827. (A) Scheme of transcriptome-guided MoA inference using Connectivity Map database (B) Frequency of matched compound’s mode of action in query (total=18) (C) Tanimoto coefficients of compounds predicted in (B), (D) Chemical structures of 5-iodotubercidin (E) Adenosine kinase activity (arbitrary unit: A.U) after LJ4827 or 5ITU treatment, Neg: Negative control, Pos: Positive control, n.s: not significant, (F) Scheme of gene ontology (Biological process: BP) analysis of downregulated genes after LJ4827 or 5ITU treatment, (G) Gene ontology (BP) analysis of downregulated genes by both LJ4827 and 5ITU treatment, (H) GSEA plot of the enrichment of the “Cell_cycle_mitotic” signature (left) and “Mitotic_spindle_Checkpoint” signature (right) in control group in comparison with treatment group (I) GSEA plot of the enrichment of the “DNA_double_strand_breeak_repair” signature in 5ITU treated group in comparison with LJ4827 treated group, (J) Immunoblotting for pH2AX at 24 hours after treatment of LJ or 5ITU (500nM) in HeLa, β-actin for equal loading control

To identify the anti-cancer MoA of LJ4827 and 5ITU against targets other than AdK, we investigated transcriptomic signatures commonly altered by LJ4827 and 5ITU treatment. We measured transcriptomic changes induced by each compound and then surveyed functions significantly associated with 3732 DEGs commonly downregulated by LJ4827 and 5ITU (Figs. 2F and S1A). Ribosome biogenesis and cell cycle-related biological processes (e.g., chromosome segregation and cell division) were significantly enriched in common DEGs (Fig. 2G). In particular, substantially downregulated genes by treatment (e.g., 5ITU or LJ4827) compared to vehicle (Cont) significantly enriched genesets of ‘mitotic spindle checkpoint’ and ‘mitotic cell cycle’ (Fig. 2H), implying that both 5ITU and LJ4827 negatively affect mitosis. Notably, compared to genes altered by LJ4827 treatment (Fig. S1B), 5ITU treatment upregulated genes associated with repair of DNA double-strand breaks (Fig. S1C), which was also revealed by geneset enrichment analysis (Fig. 2I). As predicted, a phosphorylated H2AX signal clearly was produced by 5ITU treatment (but not LJ4827), consistent with a previous study [31]. This implies that LJ4827, lacking the 5’-hydroxyl group of 4’-thioadenosine, unlike 5ITU (Fig. 2D), may impede mitosis without genotoxicity.

### LJ4827 as a putative HASPIN inhibitor

According to the assumption that chemical modification of the 5’-hydroxyl group of the kinase inhibitor LJ4425 to azide (i.e., in LJ4827) (Fig. 1C) retains the kinase inhibitor activity of LJ4827, we performed a KINOMEscan profiling assay (scanMAX) to determine the affinity of LJ4827 for 468 kinases (403 non-mutant and 65 mutant kinases) [32] (Fig. 3A). Out of 403 non-mutant kinases, 17 kinase hits were revealed after treatment with 100 nM of LJ4827 [i.e., selectivity score, S (10) = 0.042] (Fig. 3B). Among these 17 putative target kinases, HASPIN, a mitotic kinase that phosphorylates Histone H3 from late G2 throughout anaphase in mitosis [7, 8] was included (Fig. 3B). Only the four kinases CLK2, DYRK1A (in the GMGC group), MEK5 (in the STE group), and HASPIN (in the OTHER group) were present at less than 1% of the control cutoff value (Fig. 3C). Compared to IRAK4, the sixth kinase hit (Fig. 3B), the Ki value of LJ4827 toward HASPIN was 0.46 nM. This was the greatest affinity to HASPIN among the three 4’-thioadenosine analogues (Fig. S2). For further validation, a HASPIN kinase assay performed with Histone H3 peptide, which is the endogenous substrate of HASPIN, revealed a 0.155 nM IC_50_ of LJ4827 toward HASPIN (Fig. 3E). Despite similar Ki values between LJ4827 and LJ4760 (Fig. S2), LJ4827 was likely more potent than LJ4760 for inhibition of HASPIN kinase activity toward Histone H3 (Fig. 3F).

**Figure. 3.**
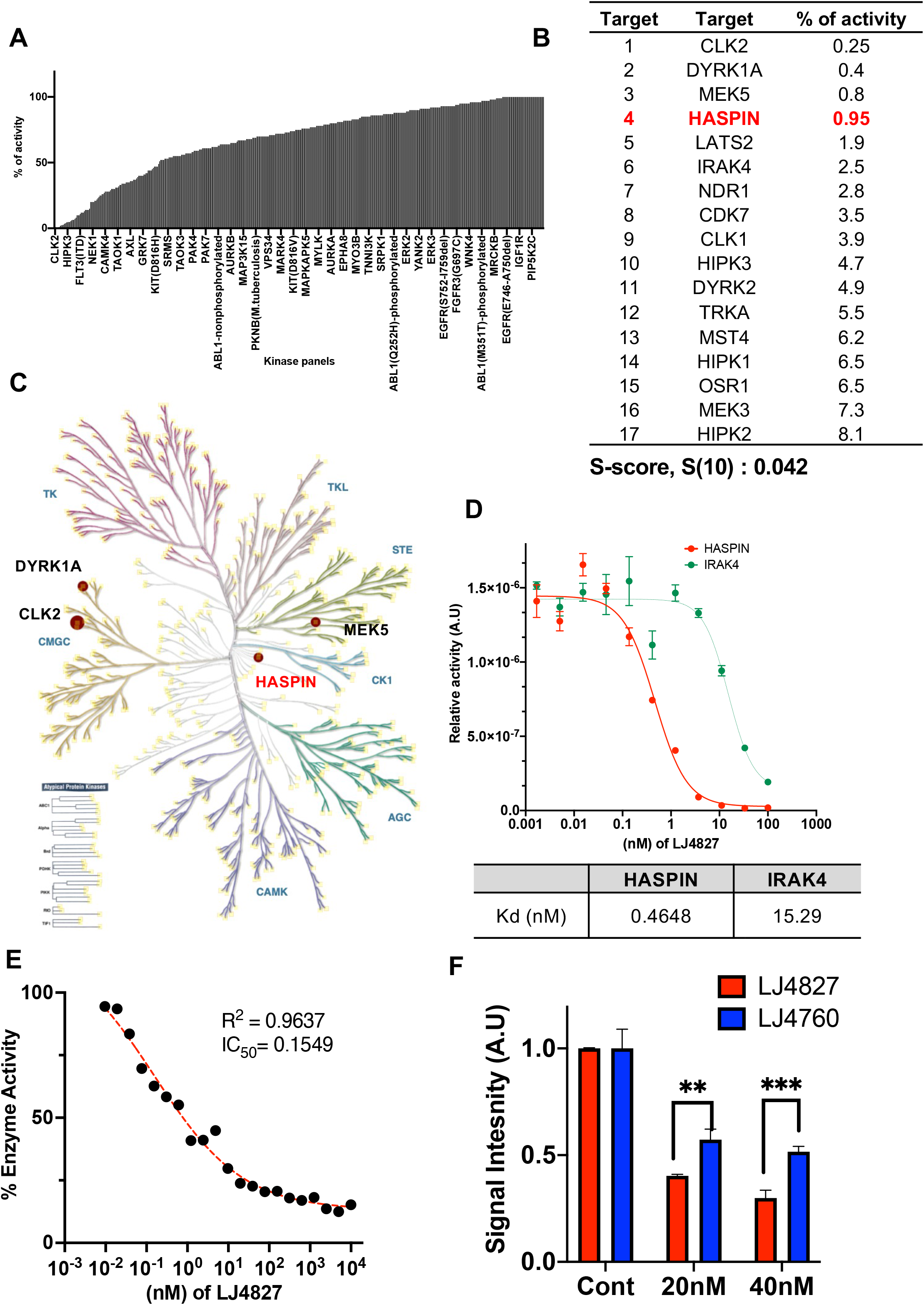
LJ4827 as a putative HASPIN inhibitor. (A) Graphical presentation of Kinomescan profiling of 100nM of LJ4827 on total 468 kinases (B) List of Hits of 100nM of LJ4827 at selective score 10 [S-score, S(10)] = (number of non-mutant kinases with %Ctrl <10)/(number of non-mutant kinases tested), HASPIN was shown in red. (C) KinMAP (Phylogenetic kinome tree) of LJ4827 (100nM) under 1% of control, HASPIN was shown in red. (D) Binding Constant (Kd) value of LJ4827 and LJ4760 for HASPIN and IRAK4, (E and F) in vitro kinase assay of HASPIN with the indicated concentrations of LJ4827 and LJ4760 using Histone H3 as a substrate

### Structural analyses of HASPIN in complex with LJ4827 and other HASPIN inhibitors

Crystal structures of the kinase domain of HASPIN in complex with LJ4827 or LJ4760 [with similar IC_50_ to HASPIN (Fig. S2)] were determined by X-ray crystallography at 2.70 Å and 2.18 Å resolution, respectively (Table S1). Both LJ4827 and LJ4760 were bound in the ATP binding site of HASPIN. The interaction modes of the adenine and ribose moieties of respective LJ4827 (Fig. 4A) and LJ4760 (Fig. 4B) with HASPIN were similar to those of AMP and 5ITU [17]. *N*^1^ and *N^6^* of the adenine moiety interacted with Glu606 and Gly608, respectively, in the HASPIN hinge region. The 2’- and 3’-hydroxyl groups of the ribose moiety interacted with Asp611 and Gly653, respectively (Figs. 4A and B). Interestingly, major structural differences in the interaction modes of LJ4827 (Fig. 4C) and LJ4760 (Fig. 4D) from AMP were observed for the moiety that replaced the 5’-hydroxyl group of ribose. In the structure of HASPIN in complex with AMP, the 5’-phosphate group interacts with Lys511 and Asp687. For 5ITU, the 5’-hydroxyl group forms a hydrogen bond with a bridging water molecule that forms a hydrogen bond with Asp687 [17]. In the structure of HASPIN in complex with LJ4827, the azide group did not interact with HASPIN. However, in the structure of HASPIN in complex with LJ4760, the amino group interacted with the backbone carbonyl group of Glu492. This interaction shortened the β1 beta-sheet, orienting the phenyl ring of Phe495 outward, unlike the structure of HASPIN in complex with LJ4827, AMP, and 5ITU (Figs. 4C and 4D). The electrostatic surface charge generated by APBS [33] showed that, compared to that of LJ4827 (Fig. 4E), LJ4760 creates a hole in the ATP binding region of HASPIN by tilting the phenyl ring outward (Fig. 4F). We speculate that Histone H3, the sole endogenous substrate of HASPIN kinase, enters through this hole, which might account for the less potent inhibition by LJ4760 compared to LJ4827, as shown in Figure 3F.

**Figure. 4.**
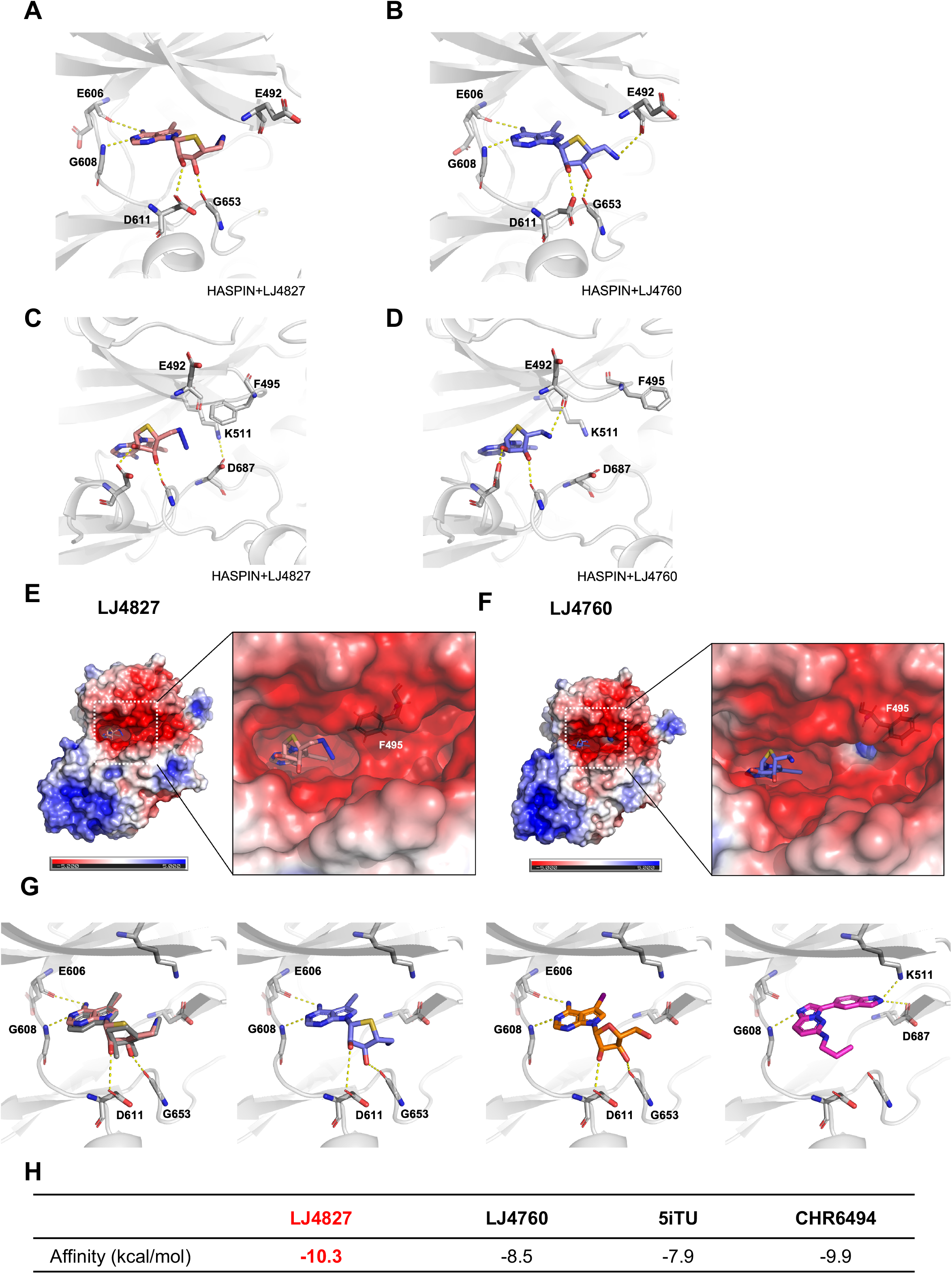
Structural analyses of HASPIN in complex with LJ4827 and other HASPIN inhibitors. (A-D) Interaction between the HASPIN hinge region and LJ4827 (A) and LJ4760 (B), Interaction between HASPIN and the azide moiety of LJ4827 (C) and the amino moiety of LJ4760 (D). (E) Surface electrostatic potential map of HASPIN in complex with LJ4827. (F) Surface electrostatic potential map of HASPIN in complex with LJ4760. (G) Predicted interaction modes between HASPIN and ligands from docking simulation. dark grey: LJ4827 from the crystal structure of HASPIN in complex with LJ4827; salmon: LJ4827 from the control docking experiment; slate blue: LJ4760; orange: 5ITU; magenta: CHR6494. Nitrogen, oxygen, sulfur, and iodine atoms are depicted as blue, red, yellow, and violet, respectively. (H) Value of binding affinity of indicated compounds on HASPIN from docking simulation

To compare the binding mode of LJ4827 with other well-characterized HASPIN inhibitors (e.g., 5ITU [34] and CHR6494 [35]), we performed a docking study for the ligands LJ4827, LJ4760, 5ITU, and CHR6494. The structure of HASPIN in complex with LJ4827 was used as a control. The interaction of LJ4827, LJ4760, and 5ITU with HASPIN was the same as mentioned above. The imidazopyridine moiety of CHR6494 interacted with Gly608, and the indazole moiety interacted with Lys511 and Asp687 (Fig. 4G). Of the four ligands tested for docking, LJ4827 showed the highest affinity (Fig. 4H).

### Retarded mitotic progression and dislocation of CPC by LJ4827

Next, in order to determine the cellular response of HASPIN inhibition by LJ4827, a HeLa cell line was used with the FUCCI (fluorescence ubiquitination cell cycle indicator) system (HeLa-FUCCI), which allows live monitoring of the cell cycle [36]. As previously described [37, 38], oscillation of green or red fluorescent signal indicates the first G2 (at the first peak of green fluorescent signal: 1^st^ G2), mitosis (at the first merge of green and red fluorescent signals: 1^st^ M), and the G1 in the second cell cycle (at the peak of red fluorescent signal: 2^nd^ G1) after release from the G1/S phase, synchronized by double thymidine block (Fig. 5A). In this setting, LJ4827 treatment markedly delayed the timing from 1^st^ G2 to 2^nd^ G1 (identified by the timing of the green and red signal peaks, respectively), similar to the response to 5ITU (Fig. 5B). In contrast, the cell cycle profile with CHR6494 indicated complete failure of mitotic progression, unlike that of LJ4827 or 5ITU (Fig. 5B). Time-lapse images of HeLa-FUCCI after compound treatment are presented (Fig. S3A and Movies S1A-D). The subsequent clonogenic assay of HeLa demonstrated that CHR6494 had the most distinct anti-proliferative activity (Fig. 5C). Nonetheless, we observed a clear pH2AX signal after CHR6494 treatment for 24 hours (Fig. 5D). According to the dissimilar cell cycle profile of CHR6494 from that of LJ4827 or 5ITU (Fig. 5B) as well as the strong pH2AX signal (Fig. 5D), we presumed that stress responses other than HASPIN inhibition (i.e., genotoxic stress) would be also responsible for the anti-proliferative activity of CHR6494. Notably, a high concentration of LJ4827 (i.e., 1.5 μM) completely inhibited mitotic entry, similar to treatment with CHR6494 (Fig. S3B and Movies S2A-D). In subsequent experiments, LJ4827 was used at no greater than 500 nM.

**Figure. 5.**
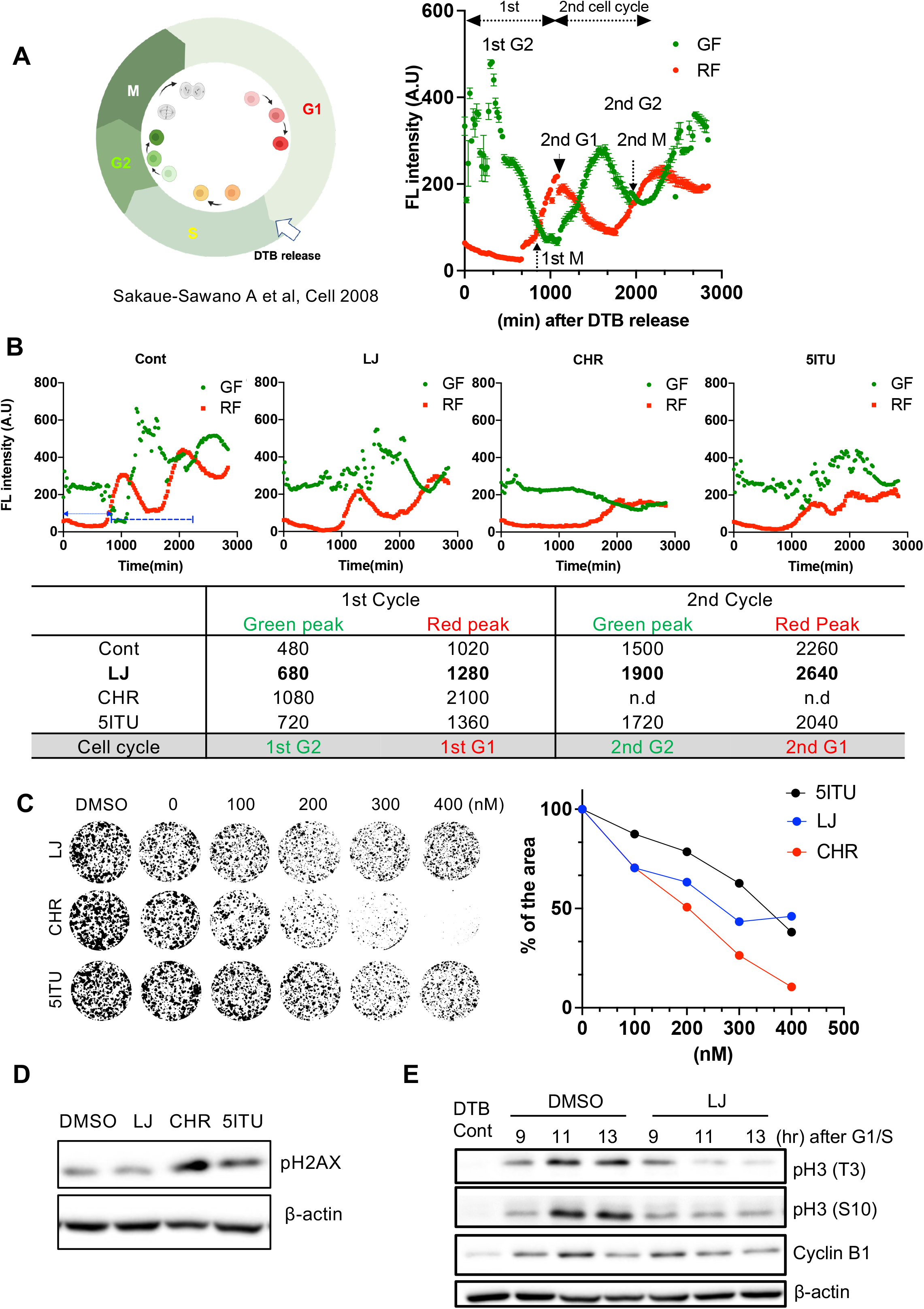
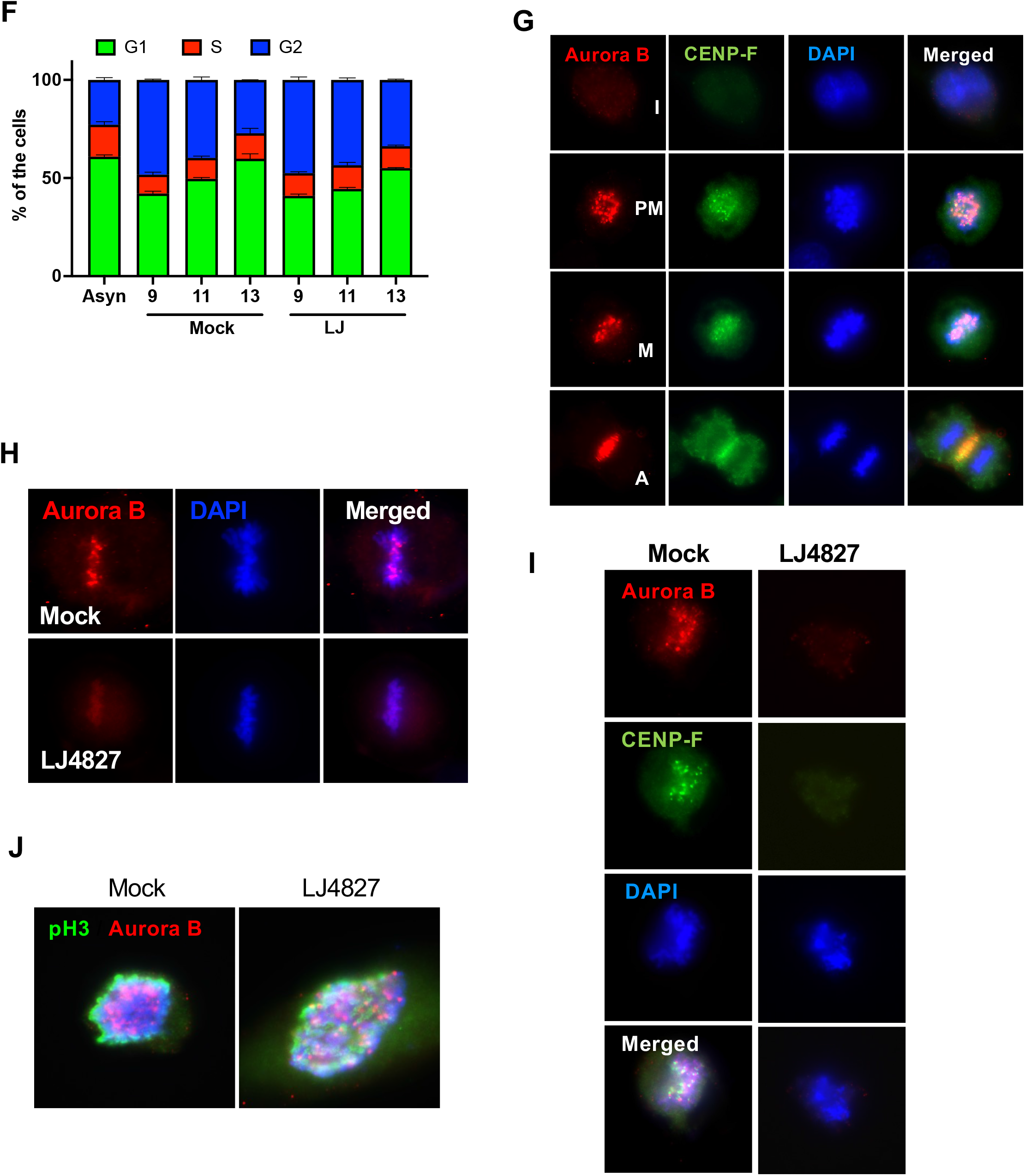
Retarded mitotic progression and dislocation of CPC by LJ4827. (A) Scheme of FUCCI system (left) and temporal intensity profiles of green (GF) or red fluorescence (RF) from FUCCI-HeLa at indicated time after release from double thymidine block (DTB) (B) temporal intensity profiles of green (GF) or red fluorescence (RF) from FUCCI-HeLa at indicated time after release from double thymidine block (DTB) with treatment of 500 nM of each compound, LJ4827: LJ, CHR6494:CHR, 5ITU respectively (top), Summary of time (minute: min) at peak of green or red fluorescence intensity after DTB, n.d: not determined (bottom) (C) Representative images of clonogenic assay after treatment of indicated dose of compounds (left), graphical quantification of relative area of survived colonies (right) (D) Immunoblotting analysis for pH2AX on A549 cells at 24 hours after 500nM treatment of each compound, β-actin for equal protein loading control (E) Immunoblotting analysis for Cyclin B1, phospho-Histone H3 [threonine 3, pH3(T3)] and phospho-Histone H3 [serine 10, pH3 (S10)] of A549 at indicative times (hr: hours) after release from G1/S (DTB Cont), synchronized by DTB in the absence (DMSO for vehicle) or presence of LJ4827 (500nM, LJ), β-actin for equal protein loading control, (F) Graphical presentation of % of cell population of G1 (green), S (red) and G2/M (blue) at indicated time after G1/S release in the absence (Mock) or presence of LJ4827 (500nM, LJ), Asyn: Asynchronized control (G) Immunofluorescent images of Aurora B (red) and CENP-F (green) in HeLa (I: interphase, PM: prometaphase, M: metaphase and A: Anaphase), (H and I) Immunofluorescent images of Aurora B (red) and CENP-F (I) in HeLa in the absence (Mock) or presence of LJ4827 (500nM, LJ) (J) Immunofluorescent images of phospho-Histone H3 (serine 10, pH3) Aurora B (red) and CENP-F (I) in hMSCs in the absence (Mock) or presence of LJ4827 (500nM, LJ)

For further validation of LJ4827 as a HASPIN inhibitor, phosphorylation of Histone H3 at threonine 3 [pH3(T3)], the sole known substrate of HASPIN during mitosis, was determined. As expected, pH3(T3) along with phosphorylated Histone H3 at serine 10 [pH3(S10)] during mitotic progression was significantly weakened by LJ4827 (Fig. 5E), which was consistent to cell cycle profile (Fig. 5F). Additionally, the pH3(T3) signal, associated with the condensed mitotic chromosome, was decreased by LJ4827 treatment (Fig. S3C). Considering that CPC recruitment at mitotic chromosome, is a downstream event of pH3(T3) after HASPIN activation, CPC location during mitosis was monitored by Aurora B, along with CENP-F, a mitotic kinetochore protein [39](Fig. 5G). Upon LJ4827 treatment, subsequent loss of pH3(T3) and location of Aurora B at the centromere (Fig. 5H) and CENP-F at the kinetochore (Fig. 5I) of the mitotic chromosome were clearly attenuated. Considering the discrete effect of genetic perturbation of HASPIN in mitosis of cancer cells [8] or normal mouse embryonic stem cells [14], we next examined the effect of HASPIN inhibition in human mesenchymal stem cells (hMSCs) as an example of a normal cell. The expression level of HASPIN in hMSCs was markedly lower than in two cancer cell lines (i.e., A549 and HeLa) (Fig. S3D). Unlike the decreased signal of Aurora B at mitotic chromosome of HeLa after LJ4827 treatment (Fig. 5H and I), that of Aurora B in hMSCs remained unaltered by LJ4827 (Fig. 5J). Accordingly, discrete effect of LJ4827 on cellular proliferation of HeLa and hMSCs was observed (Fig. S3E).

### Clinical relevance of HASPIN expression in lung cancer patients

Inspired by the observation that cancer cells exhibit higher *GSG2* expression and higher susceptibility to LJ4827 than normal cells (Fig. S2), we further investigated the relationship between *GSG2* expression and prognosis in cancer patients of the TCGA Pan-Cancer study. Likewise, expression levels of *GSG2* (Fig. 6A, paired t-test P < 0.0001) and cell cycle-related genes (Fig. 6B, FDR-adjusted p = 3.23E-48) were significantly elevated in tumor samples compared to matched normal samples. Certainly, most genes involved in mitosis (G2/M checkpoints, metaphase, and anaphase) during the cell cycle were upregulated in tumors (Figs. 6B and 6C). Considering the key role of HASPIN in mitotic progression, we speculate that high expression of *GSG2* (encoding HASPIN) allows the active mitosis required for abnormal cell proliferation in cancer. However, the expression level of *GSG2* alone would be a poor indicator for mitotic activity because *GSG2* is expressed throughout the cell cycle [7]. Accordingly, we attempted to measure the extent of active mitosis in cancer based on the expression of multiple genes associated with both HASPIN and mitosis progression. For this aim, we defined an active mitosis signature as a set of 126 mitotic genes whose expression level was both significantly elevated and correlated with *GSG2* expression in pan-cancers (Fig. 6D). Similar to the mitotic index used as a prognosis indicator by pathologists, we leveraged the active mitosis signature to score individual patients with an enrichment score through single-sample GSEA. An active mitosis signature enrichment score (AMSES) from mitosis gene set (Table S3), was significantly higher in pan-cancer (Tumor) than matched normal control (Normal) (Fig. 6E) and indicated poor prognosis in pan-cancers (Fig. S4). In similar, the clear relevance of *GSG2* expression and AMSES in lung adenocarcinoma (LUAD) patients was also determined (Fig. 6F). The AMSES well corresponded to malignancy stages of LUAD, unlike *GSG2* expression (Fig. 6G). Overall survival analysis also showed that AMSES was more indicative of LUAD prognosis than *GSG2* expression (Fig. 6H).

**Figure. 6.**
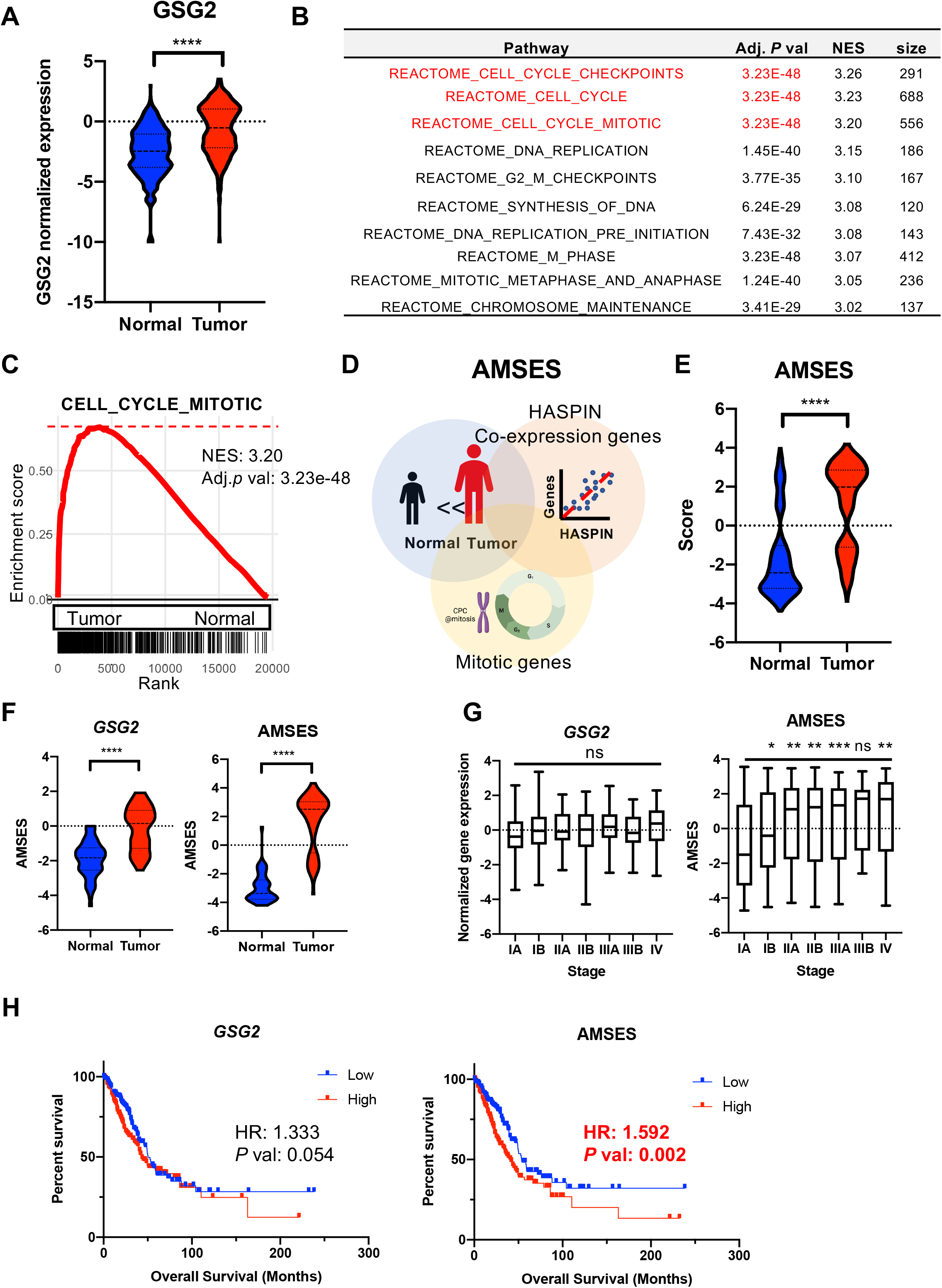
Clinical relevance of HASPIN expression in lung cancer patients. (A) Normalized *GSG2* expression in normal and tumor pairs (681 patients), (B) List of enriched REACTOME genesets in tumor group, (C) GSEA plot of the enrichment of the “Cell_cycle_mitotic” signature in tumor group in comparison with normal group, (D) Scheme for derivation of active mitosis signature enrichment score (AMSES); Selection of genes with high expression correlation with GSG2 among genes belonging to the “Cell_cycle_mitotic” gene set, (E) AMSES score in normal and tumor pairs of pan-cancer patients (681 patients), (F) Normalized GSG2 expression and AMSES score in normal and tumor pairs of LUAD patients (56 patients), (G) GSG2 expression levels by cancer stage (left), AMSES by cancer stage (right), ns: not significant (G) Kaplan-Meier survival curves of overall survival by GSG2 expression (left) and AMSES (right), HR: Hazard ratio,

### PLK1 as a synergistic partner of HASPIN inhibition

We noted that LJ4827 delayed the cell cycle but did not induce substantial cell death in the dose range of HASPIN inhibition (e.g., up to 500 nM) (Fig. S5A). To complement and potentiate anti-cancer effects of HASPIN inhibitors, synergistic targets of HASPIN inhibition have been explored by genome-wide CRISPR screening [32, 33]. Along with these efforts, an *in silico* systematic approach was recently introduced to identify synthetic lethal (SL) partner genes of cancer drugs based on a patient’s tumor transcriptome [40]. The study was performed on the basis of the assumption that the SL partner of a drug is considered a gene of which co-inhibition with the drug target(s) is associated with better prognosis in cancer patients [40]. Accordingly, we used this approach to identify synergistic partner genes of LJ4827 from the tumor transcriptome of TCGA LUAD. We hypothesized that HASPIN inactivation with a HASPIN inhibitor was epitomized by low *GSG2* expression and sought to identify genes whose low or high expression conferred survival benefits to patients. Given that HASPIN regulates chromosome behavior by interacting with other kinases during mitosis, synergistic partners were preferentially scrutinized within genes encoding kinases (i.e., readily druggable) in the predefined active mitosis signature. We first divided the patients into *GSG2*-high and *GSG2*-low groups based on *GSG2* expression and estimated the association between gene expression of each kinase and overall patient survival rates in each group (Fig. 7A). Interestingly, *BUB1B, BUB, AURKB, AURKA,* or *PLK1*, which serve as essential kinases that govern mitotic signaling for CPC regulation (Fig. 7B), showed the highest correlation to *GSG2* expression (Fig. 7C). These genes were associated with prognosis in the *GSG2*-low group (hazard ratio: HR >1) but not in the *GSG2*-high group (hazard ratio: HR <1) (Fig. 7D) and were significantly higher in tumor compared to normal cells (Fig. 7E). These findings suggest these kinases as putative druggable synergistic partners to the inhibition of HASPIN with a chemical inhibitor (i.e., LJ4827). To test this idea, a pharmacological inhibitor of each kinase was co-treated with LJ4827 to examine the synergistic effect. As shown in Figure 7F, appreciable cell death occurred by co-treatment with the PLK1 (BI2536: BI) or Aurora B (AZD1152: AZD) inhibitor. It is noteworthy that Aurora B was identified as a synergistic partner of HASPIN inhibition by genome-wide CRISPR screening [41], serving as a positive control for this approach. The strong synergistic effect of a PLK1 inhibitor (BI) with LJ4827 treatment on cytotoxicity was manifested at the 5 nM range of BI with no apparent p53 alteration (i.e., no genotoxicity) (Fig. 7G). The anti-proliferative effect of BI with LJ4827 occurred as low as the 1 nM range (Fig. 7H). According to the cell cycle profile, co-treatment of BI with LJ4827 markedly increased the Sub-G1 and 4N population (Fig. 7I), leading to increased polyploidy (Fig. 7J), suggesting that failure of proper mitosis due to simultaneous inhibition of PLK1 and HASPIN may be closely associated with cell death. The cell death induction by co-treatment occurred at the 3 nM range of BI (Fig. 7K) and 300 nM of LJ4827 (Fig. S5C). The antitumor activity of LJ4827, BI, or a combination of LJ4827 and BI was determined in a nude mouse xenograft model implanted with A549 human lung cancer cells. Compared to the vehicle control group, the tumor volumes in groups treated with LJ4827, BI, or a combination were significantly inhibited (Fig. 7L, left panel) without overt toxicity or body weight change (Fig. 7L, right panel). In turn, the tumor section from each drug regimen group was stained with TUNEL to examine cell death. Of note, a clear TUNEL-positive population was observed in the combination of LJ4827 and BI group, implying exclusive cytotoxicity toward the tumor by cotreatment with LJ4827 and BI *in vivo*.

**Figure. 7.**
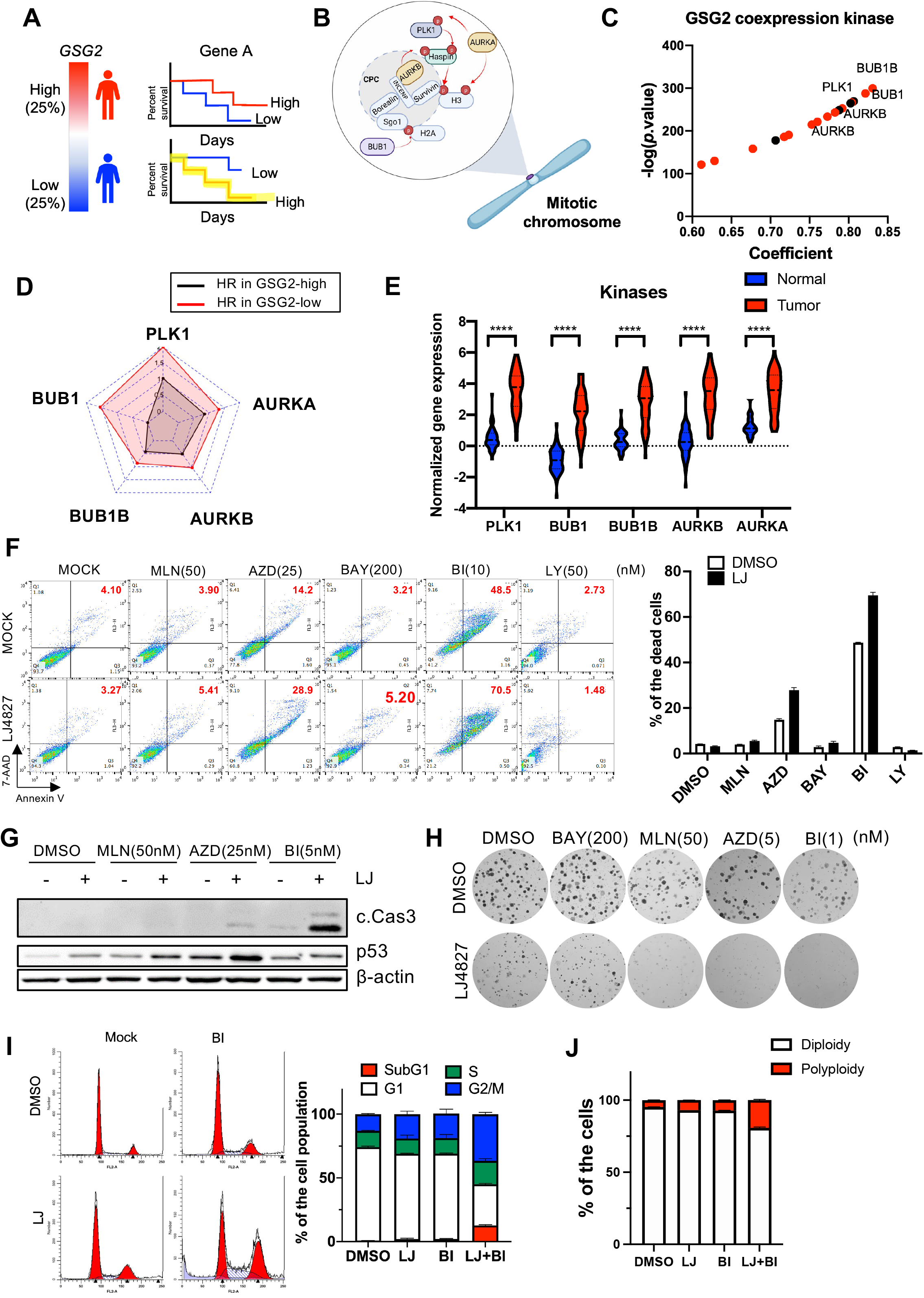
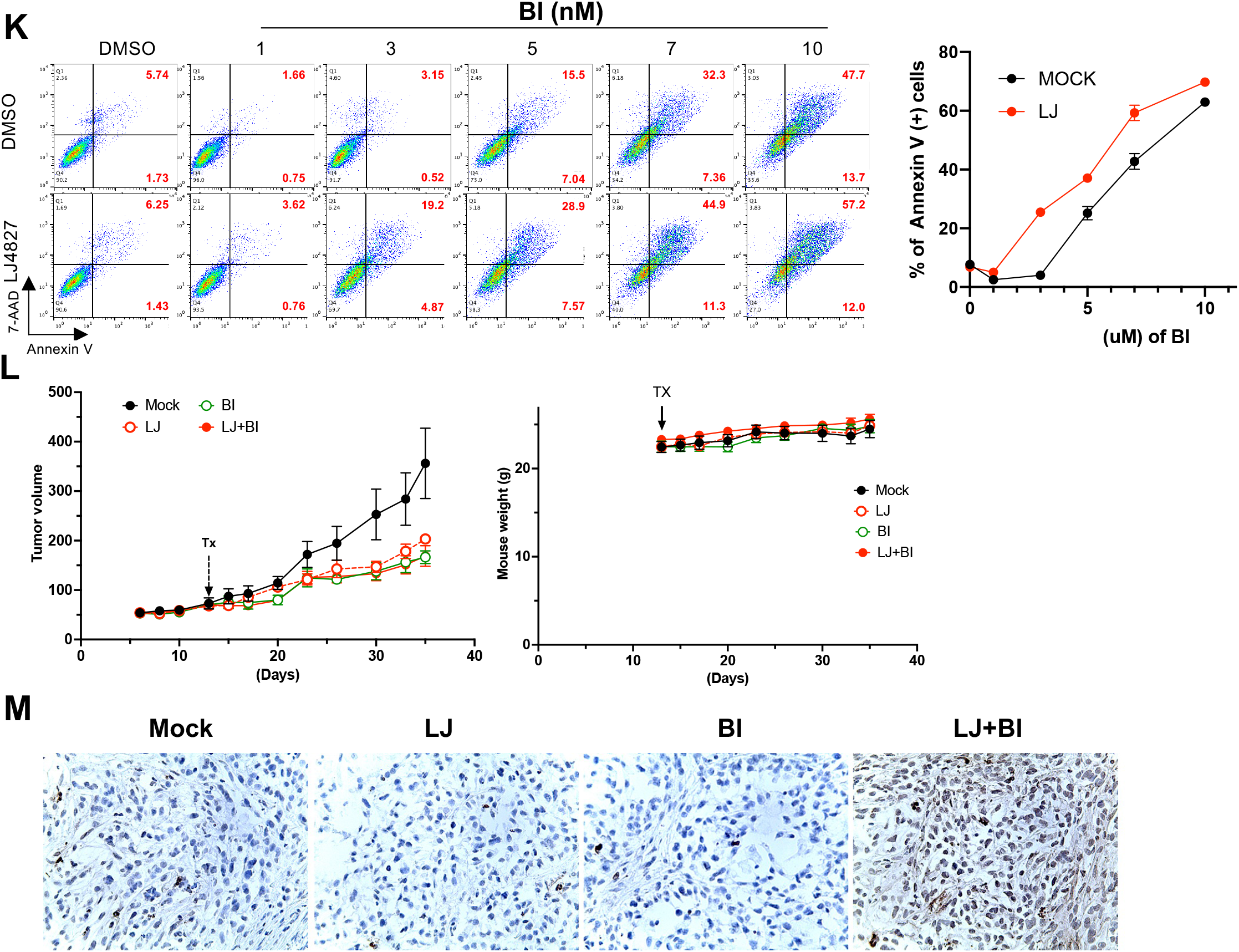
PLK1 as a synergistic partner of HASPIN inhibition. (A) Scheme for derivation synthetic lethal partner based on GSG2 expression (B) Graphical presentation of selected kinases in CPC regulation at the mitotic chromosome (C) Graph of genes (kinase shown as red), highly correlated to *GSG2* expression (D) Radar plot for hazard ratio (HR) of indicated kinase in GSG2 high (black line) or low (red line) patient group (E) Normalized gene expression of *BUB1B, BUB1, PLK1, AURKB* and *AURKA* in normal and tumor group, (F) Flow cytometry for Annexin V and 7-AAD at 48 h after treatment of indicated dose of inhibitor in A549 cells with 500nM of LJ4827 (left), MLN8237 (MLN: Aurora A/B inhibitor), AZD1152 (AZD: Aurora B inhibitor), BAY1816032 (BAY: BUB1 inhibitor), BI2536 (BI: PLK1 inhibitor) and LY3295668 (LY: Aurora A inhibitor), cell population of double positive cells for Annexin V and 7-AAD shown in red, graphical quantification of dead cells (right), (G) Immunoblotting for cleaved caspase 3 (c.Cas3) and p53 at 24 hours after indicated dose of inhibitor with 500nM of LJ4827 in A549 cells, β-actin for equal protein loading (H) Representative images of clonogenic assay after indicated dose of inhibitor with 500nM of LJ4827 in A549 cells (I) Cell cycle profile of A549 cells after treatment of BI2536 (BI: 5nM) in the absence (DMSO) or presence of LJ4827 (LJ: 500nM) (left), graphical presentation of % of the cells of subG1 (red), G1 (white), S (green) and G2/M (blue) (J) graphical presentation of % of cell population of polyploidy (red) and diploidy (while) in A549 cells at 24 hours after indicated treatment, (K) Flow cytometry for Annexin V and 7-AAD at 48hours after treatment of indicated dose of BI2536 (BI) in the absence (DMSO) or presence of LJ4827 (LJ: 500nM) (left), % of the annexin V positive population was shown in red, graphical presentation of annexin V positive cells (right), (L) Tumor volume of tumor-bearing mice after treatment (left) and changes in body weight of tumor-bearing mice after treatment (right), Tx: time at compound treatment (M) Microscopic images of TUNEL positive population of tumor ection in each treatment

## Discussion

Traditional cell-based screening for identification of anti-cancer compounds has been used as a standardized platform (e.g., NCI-60 panel) [42]. Even after identification of potential hit compounds with desirable effect, determination of MoA of the particular hit is a time-consuming but essential process for further drug development [43]. Recently, advanced computational approaches based on large datasets (i.e., chemical and biological datasets) have been proposed to facilitate this step [12, 20, 21]. Through similarity analysis based on transcriptome profile, chemical fingerprinting (i.e., https://clue.io/) and assays of chemical similarity (Fig. 2), followed by kinomescan profiling of 468 kinases (Fig. 3) and X-ray crystallography (Fig. 4), we identified HASPIN as a direct target of LJ4827, chemically modified from the 4’thio-adenosine-like multi-kinase inhibitor LJ4425 (Fig. 1). Inhibition of HASPIN with LJ4827 revealed a clear anti-mitotic effect on cancer cell lines with high expression of HASPIN by interfering with Aurora B recruitment at the mitotic centromere (Fig. 5), which would not occur in normal cells (e.g., hMSCs, Figs. 5J and S3D). We noticed that HASPIN inhibition without generation of genotoxic stress, unlike other HASPIN inhibitors (i.e., CHR6494), failed to induce massive cell death (Fig. S5A), which would limit the potential application of the HASPIN inhibitor as an anti-cancer therapeutic. For this reason, we further examined potential synergistic partners to produce cytotoxicity together with HASPIN inhibition.

Instead of genome-wide CRISPR screening, we performed *in silico* analysis in which synergistic partners of LJ4827 were screened based on the tumor transcriptome of LUAD patients. Based on ideas of a previous study [40], we sought to find synergistic partners as genes whose co-inhibition with HASPIN was associated with a good prognosis in cancer patients. We preferentially examined readily druggable (i.e., chemically inhibitable) mitotic kinases whose elevated expression represents high demand for active mitosis in cancers. Five mitotic kinases (BUB1B, BUB, AURKB, PLK1, and AURKA) closely involved in mitotic CPC regulation revealed high HR values in the HASPIN low patient group, cases that would correspond to pharmacological inhibition of HASPIN activity.

Through further biochemical analysis in lung cancer cell models, PLK1 was determined to be a potential synergistic partner of HASPIN expression. Cotreatment with BI2536, a PLK1 inhibitor in phase II clinical trials (NCT00706498 and NCT00710710), with LJ4827 induced distinct cell death at a BI2536 dose as low as 3 nM (Fig. 7). The clear synergistic effects of BI2536 and LJ4827 on cell death and mitosis would expand the application of BI2536, a first-generation PLK1 inhibitor that is no longer used in monotherapy [44]. Due to substantial toxicities in cells of actively renewing normal tissues, inhibitors of essential mitotic kinases such as PLK1, Aurora A/B, and CDK1 have not been clinically approved despite numerous studies with preclinical promise [45]. The relatively low toxicity of normal tissues of ‘target therapeutics’ results from high dependency of such ‘targets’ on cancer survival, a mechanism referred to as ‘oncogenic addiction’ [46]. In contrast, high dependency of normal cells on these essential mitotic kinases is readily evidenced by severe phenotypes after knockout [10–12], which may account for severe toxicities. In this regard, HASPIN is a promising mitotic target to potentially assure the safety of normal tissue even after complete inhibition, as neither phenotypic abnormality occurs in the HASPIN knockout mouse [13] or mESCs [14].

## Conclusion

We developed a novel genotoxicity-free HASPIN inhibitor and identified PLK1 as a synergistic partner by computational analysis, followed by *in vitro* and *in vivo* validation of the dual administration regimen.

## List of Abbreviations

HASPIN: Haploid Germ Cell□Specific Nuclear Protein Kinase
GSG2: Germ cell-specific gene 2 protein
CPC: Chromosome passenger complex
MoA: Mode of action
SMILES: Simplified Molecular Input Line Entry System
AdK: Adenosine kinase
5ITU: 5-iodotubercidin
FUCCI: Fluorescence ubiquitination cell cycle indicator
hMSCs: Human mesenchymal stem cells
AMSES: Active mitosis signature enrichment score
TCGA: The cancer genome atlas
CMap: Connectivity Map

## Declarations

### Ethics approval and consent to participate

Not applicable

### Consent to publication

Not applicable

### Availability of data and materials

The datasets used and/or analyzed during the current study are available from the corresponding author on reasonable request.

### Authors’ contributions

HJ.C and LS.J conceived the overall study design and led the experiments. EJ.K and K.M mainly conducted the experiments and data analysis, and critical discussion of the results. Y.S performed *in vitro* efficacy study. K.S synthesized the chemical compounds. J.S, SW.C and BW.H performed X-ray crystallography. SC. J and SK. L performed *in vivo* efficacy study. H.L conducted *in silico* data analysis. All authors contributed to manuscript writing and revising and endorsed the final manuscript.

## References

1. Cree IA, Tan PH, Travis WD, Wesseling P, Yagi Y, White VA, Lokuhetty D, Scolyer RA: Counting mitoses: SI(ze) matters! Mod Pathol 2021, 34:1651–1657.

2. Shapiro GI, Harper JW: Anticancer drug targets: cell cycle and checkpoint control. J Clin Invest 1999, 104:1645–1653.

3. Mills CC, Kolb EA, Sampson VB: Development of Chemotherapy with Cell-Cycle Inhibitors for Adult and Pediatric Cancer Therapy. Cancer Res 2018, 78:320–325.

4. Otto T, Sicinski P: Cell cycle proteins as promising targets in cancer therapy. Nat Rev Cancer 2017, 17:93–115.

5. Matthews HK, Bertoli C, de Bruin RAM: Cell cycle control in cancer. Nat Rev Mol Cell Biol 2022, 23:74–88.

6. Dominguez-Brauer C, Thu KL, Mason JM, Blaser H, Bray MR, Mak TW: Targeting Mitosis in Cancer: Emerging Strategies. Mol Cell 2015, 60:524–536.

7. Dai J, Sultan S, Taylor SS, Higgins JM: The kinase haspin is required for mitotic histone H3 Thr 3 phosphorylation and normal metaphase chromosome alignment. Genes Dev 2005, 19:472–488.

8. Wang F, Dai J, Daum JR, Niedzialkowska E, Banerjee B, Stukenberg PT, Gorbsky GJ, Higgins JM: Histone H3 Thr-3 phosphorylation by Haspin positions Aurora B at centromeres in mitosis. Science 2010, 330:231–235.

9. Wang F, Ulyanova NP, van der Waal MS, Patnaik D, Lens SM, Higgins JM: A positive feedback loop involving Haspin and Aurora B promotes CPC accumulation at centromeres in mitosis. Curr Biol 2011, 21:1061–1069.

10. Diril MK, Ratnacaram CK, Padmakumar VC, Du T, Wasser M, Coppola V, Tessarollo L, Kaldis P: Cyclin-dependent kinase 1 (Cdk1) is essential for cell division and suppression of DNA re-replication but not for liver regeneration. Proc Natl Acad Sci U S A 2012, 109:3826–3831.

11. Kufareva I, Abagyan R: Type-II kinase inhibitor docking, screening, and profiling using modified structures of active kinase states. J Med Chem 2008, 51:7921–7932.

12. Campillos M, Kuhn M, Gavin AC, Jensen LJ, Bork P: Drug target identification using side-effect similarity. Science 2008, 321:263–266.

13. Shimada M, Goshima T, Matsuo H, Johmura Y, Haruta M, Murata K, Tanaka H, Ikawa M, Nakanishi K, Nakanishi M: Essential role of autoactivation circuitry on Aurora B-mediated H2AX-pS121 in mitosis. Nat Commun 2016, 7:12059.

14. Soupsana K, Karanika E, Kiosse F, Christogianni A, Sfikas Y, Topalis P, Batistatou A, Kanaki Z, Klinakis A, Politou AS, Georgatos S: Distinct roles of haspin in stem cell division and male gametogenesis. Sci Rep 2021, 11:19901.

15. Amoussou NG, Bigot A, Roussakis C, Robert JH: Haspin: a promising target for the design of inhibitors as potent anticancer drugs. Drug Discov Today 2018, 23:409–415.

16. Attwood MM, Fabbro D, Sokolov AV, Knapp S, Schioth HB: Trends in kinase drug discovery: targets, indications and inhibitor design. Nat Rev Drug Discov 2021, 20:839–861.

17. Eswaran J, Patnaik D, Filippakopoulos P, Wang F, Stein RL, Murray JW, Higgins JM, Knapp S: Structure and functional characterization of the atypical human kinase haspin. Proc Natl Acad Sci U S A 2009, 106:20198–20203.

18. Elie J, Feizbakhsh O, Desban N, Josselin B, Baratte B, Bescond A, Duez J, Fant X, Bach S, Marie D, et al: Design of new disubstituted imidazo[1,2-b]pyridazine derivatives as selective Haspin inhibitors. Synthesis, binding mode and anticancer biological evaluation. J Enzyme Inhib Med Chem 2020, 35:1840–1853.

19. Terstappen GC, Schlupen C, Raggiaschi R, Gaviraghi G: Target deconvolution strategies in drug discovery. Nat Rev Drug Discov 2007, 6:891–903.

20. Lamb J: The Connectivity Map: a new tool for biomedical research. Nat Rev Cancer 2007, 7:54–60.

21. Li J, Zhu X, Chen JY: Building disease-specific drug-protein connectivity maps from molecular interaction networks and PubMed abstracts. PLoS Comput Biol 2009, 5:e1000450.

22. Iorio F, Bosotti R, Scacheri E, Belcastro V, Mithbaokar P, Ferriero R, Murino L, Tagliaferri R, Brunetti-Pierri N, Isacchi A, di Bernardo D: Discovery of drug mode of action and drug repositioning from transcriptional responses. Proc Natl Acad Sci U S A 2010, 107:14621–14626.

23. Mashelkar KK, Byun WS, Ko H, Sung K, Tripathi SK, An S, Yum YA, Kwon JY, Kim M, Kim G, et al: Discovery of a Novel Template, 7-Substituted 7-Deaza-4′-Thioadenosine Derivatives as Multi-Kinase Inhibitors. Pharmaceuticals (Basel) 2021, 14.

24. Otwinowski Z, Minor W: Processing of X-ray diffraction data collected in oscillation mode. Methods Enzymol 1997, 276:307–326.

25. Villa F, Capasso P, Tortorici M, Forneris F, de Marco A, Mattevi A, Musacchio A: Crystal structure of the catalytic domain of Haspin, an atypical kinase implicated in chromatin organization. Proc Natl Acad Sci U S A 2009, 106:20204–20209.

26. Liebschner D, Afonine PV, Baker ML, Bunkoczi G, Chen VB, Croll TI, Hintze B, Hung LW, Jain S, McCoy AJ, et al: Macromolecular structure determination using X-rays, neutrons and electrons: recent developments in Phenix. Acta Crystallogr D Struct Biol 2019, 75:861–877.

27. Emsley P, Lohkamp B, Scott WG, Cowtan K: Features and development of Coot. Acta Crystallogr D Biol Crystallogr 2010, 66:486–501.

28. Trott O, Olson AJ: AutoDock Vina: improving the speed and accuracy of docking with a new scoring function, efficient optimization, and multithreading. J Comput Chem 2010, 31:455–461.

29. Schuttelkopf AW, van Aalten DM: PRODRG: a tool for high-throughput crystallography of protein-ligand complexes. Acta Crystallogr D Biol Crystallogr 2004, 60:1355–1363.

30. Martin YC, Kofron JL, Traphagen LM: Do structurally similar molecules have similar biological activity? J Med Chem 2002, 45:4350–4358.

31. Zhang X, Jia D, Liu H, Zhu N, Zhang W, Feng J, Yin J, Hao B, Cui D, Deng Y, et al: Identification of 5-Iodotubercidin as a genotoxic drug with anti-cancer potential. PLoS One 2013, 8:e62527.

32. Fabian MA, Biggs WH, 3rd, Treiber DK, Atteridge CE, Azimioara MD, Benedetti MG, Carter TA, Ciceri P, Edeen PT, Floyd M, et al: A small molecule-kinase interaction map for clinical kinase inhibitors. Nat Biotechnol 2005, 23:329–336.

33. Jurrus E, Engel D, Star K, Monson K, Brandi J, Felberg LE, Brookes DH, Wilson L, Chen J, Liles K, et al: Improvements to the APBS biomolecular solvation software suite. Protein Sci 2018, 27:112–128.

34. De Antoni A, Maffini S, Knapp S, Musacchio A, Santaguida S: A small-molecule inhibitor of Haspin alters the kinetochore functions of Aurora B. J Cell Biol 2012, 199:269–284.

35. Huertas D, Soler M, Moreto J, Villanueva A, Martinez A, Vidal A, Charlton M, Moffat D, Patel S, McDermott J, et al: Antitumor activity of a small-molecule inhibitor of the histone kinase Haspin. Oncogene 2012, 31:1408–1418.

36. Sakaue-Sawano A, Kurokawa H, Morimura T, Hanyu A, Hama H, Osawa H, Kashiwagi S, Fukami K, Miyata T, Miyoshi H, et al: Visualizing spatiotemporal dynamics of multicellular cell-cycle progression. Cell 2008, 132:487–498.

37. Go YH, Lee HJ, Kong HJ, Jeong HC, Lee DY, Hong SK, Sung SH, Kwon OS, Cha HJ: Screening of cytotoxic or cytostatic flavonoids with quantitative Fluorescent Ubiquitination-based Cell Cycle Indicator-based cell cycle assay. R Soc Open Sci 2018, 5:181303.

38. Koh SB, Mascalchi P, Rodriguez E, Lin Y, Jodrell DI, Richards FM, Lyons SK: A quantitative FastFUCCI assay defines cell cycle dynamics at a single-cell level. J Cell Sci 2017, 130:512–520.

39. Liao H, Winkfein RJ, Mack G, Rattner JB, Yen TJ: CENP-F is a protein of the nuclear matrix that assembles onto kinetochores at late G2 and is rapidly degraded after mitosis. J Cell Biol 1995, 130:507–518.

40. Lee JS, Nair NU, Dinstag G, Chapman L, Chung Y, Wang K, Sinha S, Cha H, Kim D, Schperberg AV, et al: Synthetic lethality-mediated precision oncology via the tumor transcriptome. Cell 2021, 184:2487–2502 e2413.

41. Huang M, Feng X, Su D, Wang G, Wang C, Tang M, Paulucci-Holthauzen A, Hart T, Chen J: Genome-wide CRISPR screen uncovers a synergistic effect of combining Haspin and Aurora kinase B inhibition. Oncogene 2020, 39:4312–4322.

42. Sharma SV, Haber DA, Settleman J: Cell line-based platforms to evaluate the therapeutic efficacy of candidate anticancer agents. Nat Rev Cancer 2010, 10:241–253.

43. Schenone M, Dancik V, Wagner BK, Clemons PA: Target identification and mechanism of action in chemical biology and drug discovery. Nat Chem Biol 2013, 9:232–240.

44. Gutteridge RE, Ndiaye MA, Liu X, Ahmad N: Plk1 Inhibitors in Cancer Therapy: From Laboratory to Clinics. Mol Cancer Ther 2016, 15:1427–1435.

45. Yan VC, Butterfield HE, Poral AH, Yan MJ, Yang KL, Pham CD, Muller FL: Why Great Mitotic Inhibitors Make Poor Cancer Drugs. Trends Cancer 2020, 6:924–941.

46. Sharma SV, Settleman J: Oncogene addiction: setting the stage for molecularly targeted cancer therapy. Genes Dev 2007, 21:3214–3231.

