## Supplemental Files for "Identification of a novel HASPIN inhibitor and its synergism with the PLK1 inhibitor"

This PDF file includes:

Supplemental Scheme S1

Supplemental Materials and Methods

Supplemental Figure legends

Supplementary Figures 1-5

#### Supplemental Materials and Methods

#### Chemical Synthesis of Final Compounds 4, 5, and 8 (Scheme S1)

**Scheme S1**

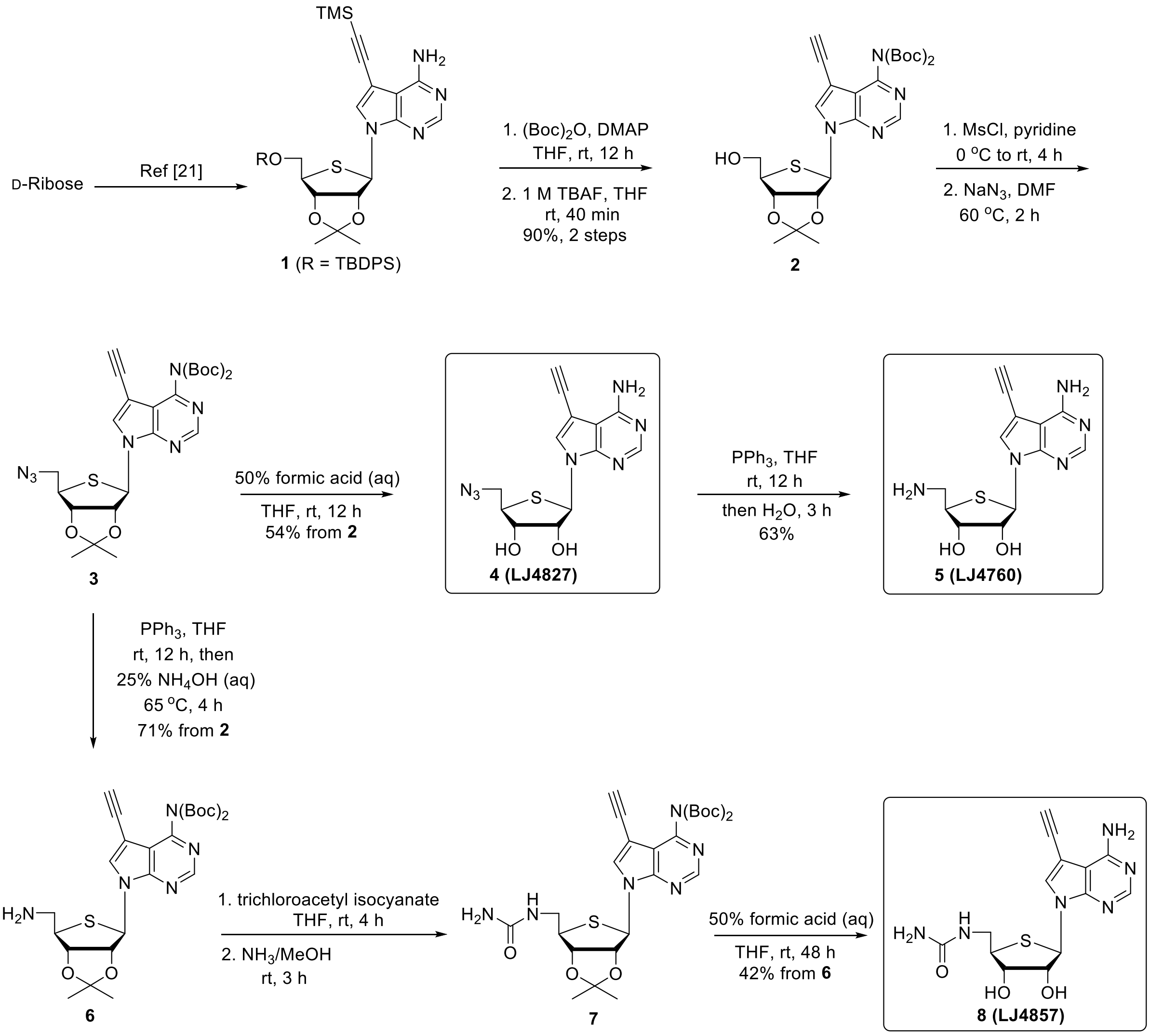


#### General Methods

Proton ($^1$H) and carbon ($^{13}$C) nuclear magnetic resonance spectra were recorded on a JEOL JNM-GCX (400/100 MHz) or Bruker AMX-500 (500/125 MHz) spectrometer in the solvent indicated. Chemical shifts are given in $\delta$ values, and are referenced to the solvent peak. Coupling constants ($J$) are reported in Hertz (Hz). High-resolution mass (HRMS) data were obtained on a Thermo LCQ XP instrument. UV spectra were recorded on a U-3000 made by Hitachi. Jasco III was used to measure optical rotations and $[\alpha]^{25}_{D}$ values are given in $10^{-1}$ deg $cm^2$ $g^{-1}$. Melting points were recorded on a Barnstead electrothermal 9100 instrument and are uncorrected. The TLC spots analyzed under ultraviolet light at 254 nm were further visualized by *p*-anisaldehyde or phosphomolybdic acid stain solution. Column chromatography was performed on silica gel (Kieselgel 60, 70-230 mesh, Merck).

***tert*-Butyl (*tert*-butoxycarbonyl)(5-ethynyl-7-((3a*R*,4*R*,6*R*,6a*S*)-6-(hydroxymethyl)-2,2-dimethyltetrahydrothieno[3,4-*d*][1,3]dioxol-4-yl)-7*H*-pyrrolo[2,3-*d*]pyrimidin-4-yl)carbamate (2)**. To a solution of **1** [21] (0.85 g, 1.29 mmol) in THF (8.5 mL) was added DMAP (0.47 g, 3.88 mmol) followed by di-*tert*-butyl dicarbonate (1.77 mL, 7.76 mmol) at room temperature. After stirring at same temperature for 12 h, the volatiles were evaporated and the residue was partitioned between water (300 mL) and ethyl acetate (3 × 200 mL). The organic layer was separated, combined, dried over $MgSO_4$, filtered, and concentrated to give the crude compound.

This crude compound was dissolved in anhydrous THF (5.8 mL) and a solution of 1 M TBAF in THF (2.64 mL, 2.64 mmol) was added to it under nitrogen atmosphere and the resulting reaction mixture was stirred at room temperature for 40 min. The reaction was quenched with saturated aqueous $NH_4Cl$ solution (5 mL) and extracted with ethyl acetate (3 × 150 mL). The combined organic layer was washed with brine (100 mL), dried over $MgSO_4$, filtered, and concentrated to give the residue which on purification afforded **2** (0.63 g, 90% in 2 steps) as sticky mass; silica gel column chromatography ($CH_2Cl_2$/MeOH, 49:1); $[\alpha]^{25}_{D}$ -74.61 (*c* 5.65, $CH_3OH$); UV ($CH_3OH$) $\lambda_{max}$ 235.29 nm; $^1H$ NMR ($CD_3OD$, 400 MHz): $\delta$ 8.78 (s, 1H), 8.30 (s, 1H), 6.46 (d, $J$ = 2.8 Hz, 1H), 5.21 (dd, $J$ = 5.6, 3.2 Hz, 1H), 5.02 (dd, $J$ = 5.2, 1.2 Hz, 1H), 3.79-3.74 (m, 3H), 3.67 (s, 1H), 1.61 (s, 3H), 1.34 (s, 21H); $^{13}C$ NMR ($CD_3OD$, 100 MHz): $\delta$

152.1, 151.6, 151.4, 149.9, 133.1, 116.6, 111.9, 95.8, 89.3, 85.6, 83.7, 81.1, 74.3, 67.4, 63.7, 55.7, 26.7, 26.5, 24.1; HRMS (ESI-Q-TOF) *m*/*z* [M + H]$^+$ for $C_{26}H_{35}N_4O_7S$ calculated 547.2221, found 547.2214.

***tert*-Butyl (7-((3a*R*,4*R*,6*R*,6a*S*)-6-(azidomethyl)-2,2-dimethyltetrahydrothieno[3,4-*d*][1,3]dioxol-4-yl)-5-ethynyl-7*H*-pyrrolo[2,3-*d*]pyrimidin-4-yl)(*tert*-butoxycarbonyl) carbamate (3)**. Mesyl chloride (0.1 mL, 1.46 mmol) was added to a solution of **2** (0.40 g, 0.73 mmol) in anhydrous pyridine (15 mL/mmol) at 0 °C under nitrogen atmosphere and the reaction mixture was stirred at room temperature for 4 h. The volatiles were evaporated under vacuum and the residue was partitioned between water (220 mL) and ethyl acetate (3 × 160 mL). The separated organic layers were combined, dried over $MgSO_4$, filtered, and evaporated under vacuum below 25 °C water bath temperature. The crude mesylate was dissolved in anhydrous DMF (5.5 mL/mmol) and sodium azide (0.14 g, 2.16 mmol) was added portion wise at room temperature. The reaction mixture was transferred to preheated bath at 60 °C and stirred at the same temperature for 2 h. The solvent was evaporated under reduced pressure and the residue was partitioned between water (200 mL) and ethyl acetate (3 × 180 mL). The combined organic layers were dried over $MgSO_4$, filtered and concentrated to give crude **3** ($R_f$ = 0.50, TLC eluent = hexane/ethyl acetate, 4:1) which was used as in the next step; HRMS (ESI-Q-TOF) *m*/*z* [M + H]$^+$ for $C_{26}H_{34}N_7O_6S$ calculated 572.2286, found 572.2277.

**(2*R*,3*R*,4*S*,5*R*)-2-(4-amino-5-ethynyl-7*H*-pyrrolo[2,3-*d*]pyrimidin-7-yl)-5-(azidomethyl)tetrahydrothiophene-3,4-diol (4).** To the crude intermediate **3** was added 50% aqueous formic acid solution (27 mL) and THF (5 mL). The resulting solution was stirred at room temperature for 12 h. Acidic solution was basified using a weakly basic anion-exchange resin (Dowex® 66 free base) and stirred for additional 3 h, filtered, and concentrated in vacuum. The residue was purified by silica gel column chromatography ($CH_2Cl_2$/MeOH, 24:1) to obtain

**4** (0.13 g, 54% in 3 steps; 0.40 g scale of **2**) as white solid; mp 110-112 ºC; $[\alpha]^{25}_{D}$ -30.23 (*c* 2.36, $CH_3OH$); UV ($CH_3OH$) $\lambda_{max}$ 283.53 nm; IR $\nu_{max}$ strong band at 2169 $cm^{-1}$; $^1H$ NMR (DMSO-$d_6$, 500 MHz): $\delta$ 8.12 (s, 1H), 7.94 (s, 1H), 6.10 (d, $J$ = 6.8 Hz, 1H), 5.59 (d, $J$ = 6.2 Hz, 1H), 5.51 (d, $J$ = 4.5 Hz, 1H), 4.52-4.48 (m, 1H), 4.30 (s, 1H), 4.09-4.07 (m, 1H), 3.86 (dd, $J$ = 12.5, 6.9 Hz, 1H), 3.75 (dd, $J$ = 12.5, 7.3 Hz, 1H), 3.38-3.34 (m, 1H); $^{13}C$ NMR (DMSO-$d_6$, 100 MHz): $\delta$ 157.4, 152.9, 149.9, 127.4, 102.2, 94.3, 83.2, 77.2, 76.8, 73.8, 61.3, 53.7, 50.1; HRMS (ESI-Q-TOF) *m/z* $[M + H]^+$ for $C_{13}H_{14}N_7O_2S$ calculated 332.0924, found 332.0923.

**(2*R*,3*R*,4*S*,5*R*)-2-(4-amino-5-ethynyl-7*H*-pyrrolo[2,3-*d*]pyrimidin-7-yl)-5-(aminomethyl)tetrahydrothiophene-3,4-diol (5).** To a solution of **4** (0.07 g, 0.21 mmol) in THF (5.6 mL) was added triphenylphosphine (0.11 g, 0.42 mmol) and the resulting solution was stirred at room temperature for 12 h. Water was added to the reaction mixture and stirred at room temperature for another 3 h. The volatiles were evaporated and the residue was purified by silica gel column chromatography ($CH_2Cl_2$/MeOH, 89:11) to afford **5** (0.04 g, 63%) as white solid; mp 125-127 ºC; $[\alpha]^{25}_{D}$ -26.98 (*c* 0.56, $CH_3OH$); UV ($CH_3OH$) $\lambda_{max}$ 279.41 nm; $^1H$ NMR (DMSO-$d_6$, 400 MHz): $\delta$ 8.12 (s, 1H), 7.94 (s, 1H), 6.07 (d, $J$ = 6.7 Hz, 1H), 5.45 (d, $J$ = 6.1 Hz, 1H), 4.45-4.41 (m, 1H), 4.30 (s, 1H), 4.12 (merged dd, $J_1 = J_2$ = 3.6 Hz, 1H), 3.26-3.21 (m, 1H), 3.0 (dd, $J$ = 13.4, 6.1 Hz, 1H), 2.81 (dd, $J$ = 13.4, 7.3 Hz, 1H); $^{13}C$ NMR (DMSO-$d_6$, 100 MHz): $\delta$ 157.4, 152.8, 149.9, 127.6, 102.2, 94.1, 83.1, 77.4, 77.3, 73.9, 61.1, 54.0, 45.4; HRMS (ESI-Q-TOF) *m/z* $[M + H]^+$ for $C_{13}H_{16}N_5O_2S$ calculated 306.1019, found 306.1031.

***tert*-Butyl (7-((3a*R*,4*R*,6*R*,6a*S*)-6-(aminomethyl)-2,2-dimethyltetrahydrothieno[3,4-*d*][1,3]dioxol-4-yl)-5-ethynyl-7*H*-pyrrolo[2,3-*d*]pyrimidin-4-yl)(*tert*-butoxycarbonyl)carbamate (6).** The crude **3** (1 equiv) was dissolved in anhydrous THF (27 mL/mmol) and to it was added triphenylphosphine (2 equiv) under nitrogen atmosphere. The reaction mixture was stirred at room temperature for 12 h. Ammonium hydroxide (25%, 1.8

mL/mmol) was added to the resulting solution and heated to 65 °C for 4 h. The volatiles were evaporated under vacuum and the residue was purified by silica gel column chromatography ($CH_2Cl_2$/MeOH, 19:1) to afford **6** (0.35 g, 71% in 3 steps; 0.50 g scale of **2**) as pale yellow sticky mass; $[\alpha]^{25}_{D}$ -27.61 (*c* 1.52, $CH_3OH$); UV ($CH_3OH$) $\lambda_{max}$ 280.92 nm; $^1H$ NMR ($CD_3OD$, 400 MHz): $\delta$ 8.80 (s, 1H), 8.11 (s, 1H), 6.42 (d, $J$ = 3.2 Hz, 1H), 5.32 (dd, $J$ = 5.6, 2.8 Hz, 1H), 5.01 (dd, $J$ = 6.0, 2.8 Hz, 1H), 3.70-3.66 (s merged with m, 2H), 3.05-3.00 (m, 1H), 2.88-2.85 (m, 1H), 1.60 (s, 3H), 1.34 (s, 21H); HRMS (ESI-Q-TOF) *m/z* $[M + H]^+$ for $C_{26}H_{36}N_5O_6S$ calculated 546.2381, found 546.2372.

***tert*-Butyl (*tert*-butoxycarbonyl)(7-((3a*R*,4*R*,6*R*,6a*S*)-2,2-dimethyl-6-(ureidomethyl)tetrahydrothieno[3,4-*d*][1,3]dioxol-4-yl)-5-ethynyl-7*H*-pyrrolo[2,3-*d*]pyrimidin-4-yl)carbamate (7).** Trichloroacetyl isocyanate (0.075 g, 0.40 mmol) was added dropwise to a stirred solution of **6** (0.22 g, 0.40 mmol) in anhydrous THF (6.6 mL) and the reaction mixture was stirred at room temperature for 4 h. Reaction was quenched with water (25 mL) and extracted with ethyl acetate (3 × 60 mL). Organics were combined, dried, filtered and concentrated to give crude which was utilized for the next step without any purification.

To the crude was added $NH_3$/*tert*-BuOH (5 mL) and stirred at room temperature for 3 h. Volatiles were evaporated under vacuum and the crude residue **7** ($R_f$ = 0.60, TLC eluent = $CH_2Cl_2$/MeOH, 9:1) was subjected to the next reaction; HRMS (ESI-Q-TOF) *m/z* $[M + H]^+$ for $C_{27}H_{37}N_6O_7S$ calculated 589.2439, found 589.2447.

**1-(((2*R*,3*S*,4*R*,5*R*)-5-(4-amino-5-ethynyl-7*H*-pyrrolo[2,3-*d*]pyrimidin-7-yl)-3,4-dihydroxytetrahydrothiophen-2-yl)methyl)urea (8).** The crude **7** was converted to **8** (0.06 g, 42% in 3 steps) as white solid, by following the procedure described for **4**; silica gel column chromatography ($CH_2Cl_2$/MeOH, 89:11); mp 114-116 °C; $[\alpha]^{25}_{D}$ -21.79 (*c* 0.06, $CH_3OH$); UV ($CH_3OH$) $\lambda_{max}$ 281.18 nm; $^1H$ NMR ($CD_3OD$, 400 MHz): $\delta$ 8.11 (s, 1H), 7.79 (s, 1H), 6.16 (d,

$J$ = 6.0 Hz, 1H), 4.47 (dd, $J$ = 5.6, 3.6 Hz, 1H), 4.16-4.14 (m, 1H), 3.73 (s, 1H), 3.60-3.52 (m, 1H), 3.50-3.41 (m, 2H); $^{13}$C NMR (DMSO-$d_6$, 125 MHz): $\delta$ 158.5, 157.4, 152.8, 150.0, 127.4, 102.2, 94.2, 83.0, 77.3, 77.1, 73.9, 61.0, 51.8; HRMS (ESI-Q-TOF) $m/z$ $[M + H]^+$ for $C_{14}H_{17}N_6O_3S$ calculated 349.1077, found 349.1092.

**Cell culture**

Hela, A549 and hMSCs cell lines were maintained in Dulbecco's modified Eagle Medium (DMEM). DMEM were supplemented with 10% (v/v) fetal bovine serum, gentamicin (50 μg/mL) at 37 °C in a humidified atmosphere of 5% $CO_2$ in the air.

**Annexin-V & 7-AAD staining**

To determine the population of dead cells, Annexin-V & 7-AAD staining was carried out in accordance with the manufacturer's instructions (559763, BD Pharmingen). Results were analyzed using a Becton-Dickinson FACS Calibur-1 with the Cell-QUEST software.

**PI staining & Cell cycle profiling**

PI staining to determine cell cycle distribution (Sub G1, G1, S and G2/M) based on DNA content. The stained cells were analyzed using Becton-Dickinson FACS Calibur-1 and Modfit 4.1 cell cycle analysis software.

**Cell synchronization**

Using double thymidine block (DTB), FUCCI-HeLa cells were synchronized at the G1/S boundary. In detail, cells were treated with 2.5 mM thymidine for 16 h and were released back with normal medium for 8 h. After second thymidine treatment with 2.5 mM thymidine for an additional 16 h, cells were arrested at G1/S boundary. Cells were live monitored with the JULI-stage live image machine (NanoEntek).

**Time-lapse imaging and compensation**

Time-lapse images were acquired at constant time points (Bright field (BF), GFP, and RFP 3-channel. LED power: 1, Bright: 4, RFP is 7. Exposure: 400 ms, BF is 70 ms. Images captured with auto-focusing in all channels and cycles) using JULI-stage software (NanoEntek), and contrast of both green and red fluorescence compensated using batch editor of photoscape software (Auto contrast: middle. Contrast: low, +5. Erode). Attached cell counter of JULI-stat (NanoEntek) was used as analyzing software (intensity of min: 10, max: 255, counted for each pixel) [1].

**Clonogenic assay**

A549 and Hela cells were seeded in 6 well plate at cell concentrations estimated to yield 20–100 colonies/well. After 24 h of incubation, cells were pre-treated with 20 or 30 μM. Cells were cultured for 10–14 d, and colonies were counted using Image J.

**Supplemental Figure legends**

**Figure. S1** (A) Scheme of gene ontology (BP) analysis of downregulated genes after LJ4827 and 5ITU treatment, (B) Gene ontology (BP) analysis of downregulated genes by only LJ4827 treatment, (C) Gene ontology (BP) analysis of downregulated genes by only 5ITU treatment.

**Figure. S2** Binding Constant (Kd) value of LJ4827, LJ4857 and LJ4760 for HASPIN (left),

**Figure. S3** (A) Timelapse images of cell cycle after HASPIN inhibitors treatment, 1 frame = 20-minute, (B) Timelapse images of cell cycle after LJ 4827 concentration dependent treatment, (C) Fluorescent microscopic images of phospho-Histone H3 [threonine 3: PH3 (T3)] of HeLa at interphase (I) or metaphase (M) (D) Immunoblotting analysis for HASPIN on HeLa, hMSCs and A549, β-actin for equal protein loading, (E) Proliferation rate of HeLa and hMSCs after 500nM of LJ4827

**Figure. S4** Kaplan-Meier survival curves for overall survival with high AMSES in PAN-cancer

**Figure. S5** (A) Flow cytometry for Annexin V and 7-AAD at 48 hours after indicated dose of LJ4827 (left), graphical presentation of dual positive population with Annexin V and 7-AAD staining (right) (B and C) Kaplan-Meier plot for overall survival with BUB1, BUB1B, PLK1 and AURKB expression in GSG2 low (B) and high (C) patients (D) Flow cytometry for Annexin V and 7-AAD at 48 h after indicated dose of LJ 4827 concentration with and without BI2536 (left), the quantitative measurement of Annexin V and 7-AAD both negative cells (right)

**Movie S1** (A) Timelapse video of cell cycle after vehicle treatment, (B) Timelapse video of cell cycle after LJ4827 treatment, (C) Timelapse video of cell cycle after CHR6494 treatment, (D) Timelapse video of cell cycle after 5ITU treatment

**Movie S2** (A) Timelapse video of cell cycle after vehicle treatment, (B) Timelapse video of cell cycle after LJ4827 (0.5uM) treatment, (C) Timelapse video of cell cycle after LJ4827 (1uM) treatment, (D) Timelapse video of cell cycle after LJ4827 (1.5uM) treatment

**Table S1** Queried result of CMap with LJ4827 transcriptome

**Table S2** Data collection and refinement statistics

**Table S3** REACTOME geneset enrichment analysis result in tumor group compared to normal group

**Table S4** List of active mitosis gene set

#### Reference

1. Go YH, Lee HJ, Kong HJ, Jeong HC, Lee DY, Hong SK, Sung SH, Kwon OS, Cha HJ: **Screening of cytotoxic or cytostatic flavonoids with quantitative Fluorescent Ubiquitination-based Cell Cycle Indicator-based cell cycle assay.** *R Soc Open Sci* 2018, **5:**181303.

### Figure S1

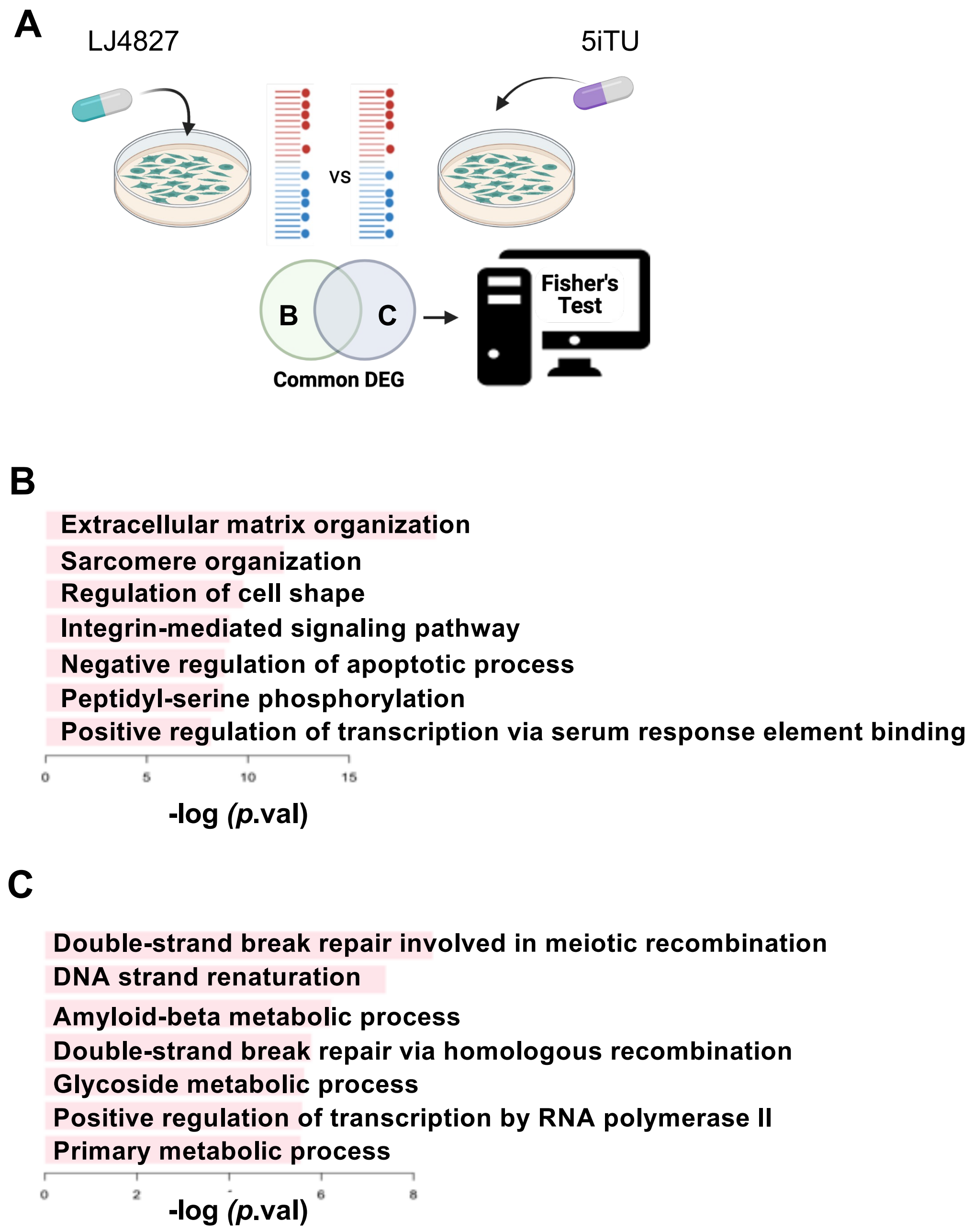
A
LJ4827
5iTU
vs
B
C
Common DEG
Fisher's Test
B
Extracellular matrix organization
Sarcomere organization
Regulation of cell shape
Integrin-mediated signaling pathway
Negative regulation of apoptotic process
Peptidyl-serine phosphorylation
Positive regulation of transcription via serum response element binding
0
5
10
15
-log (p.val)
C
Double-strand break repair involved in meiotic recombination
DNA strand renaturation
Amyloid-beta metabolic process
Double-strand break repair via homologous recombination
Glycoside metabolic process
Positive regulation of transcription by RNA polymerase II
Primary metabolic process
0
2
6
8
-log (p.val)

**Figure S2**

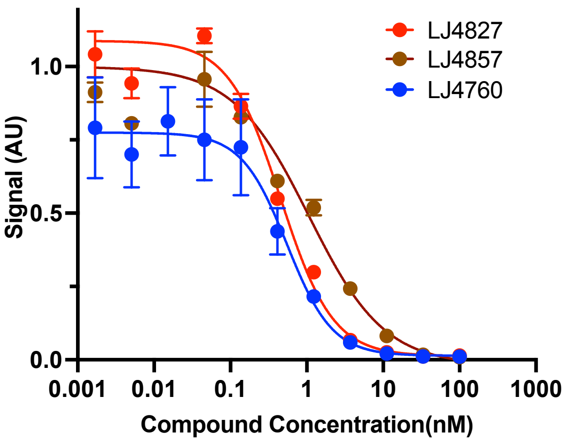


| | LJ4827 | LJ4857 | LJ4760 |
|---|---|---|---|
| Affinity (Kcal/mol) | -8.9 | -9.0 | -8.3 |
| Ki (nM) | **0.453** | 1.065 | 0.548 |

#### Figure S3

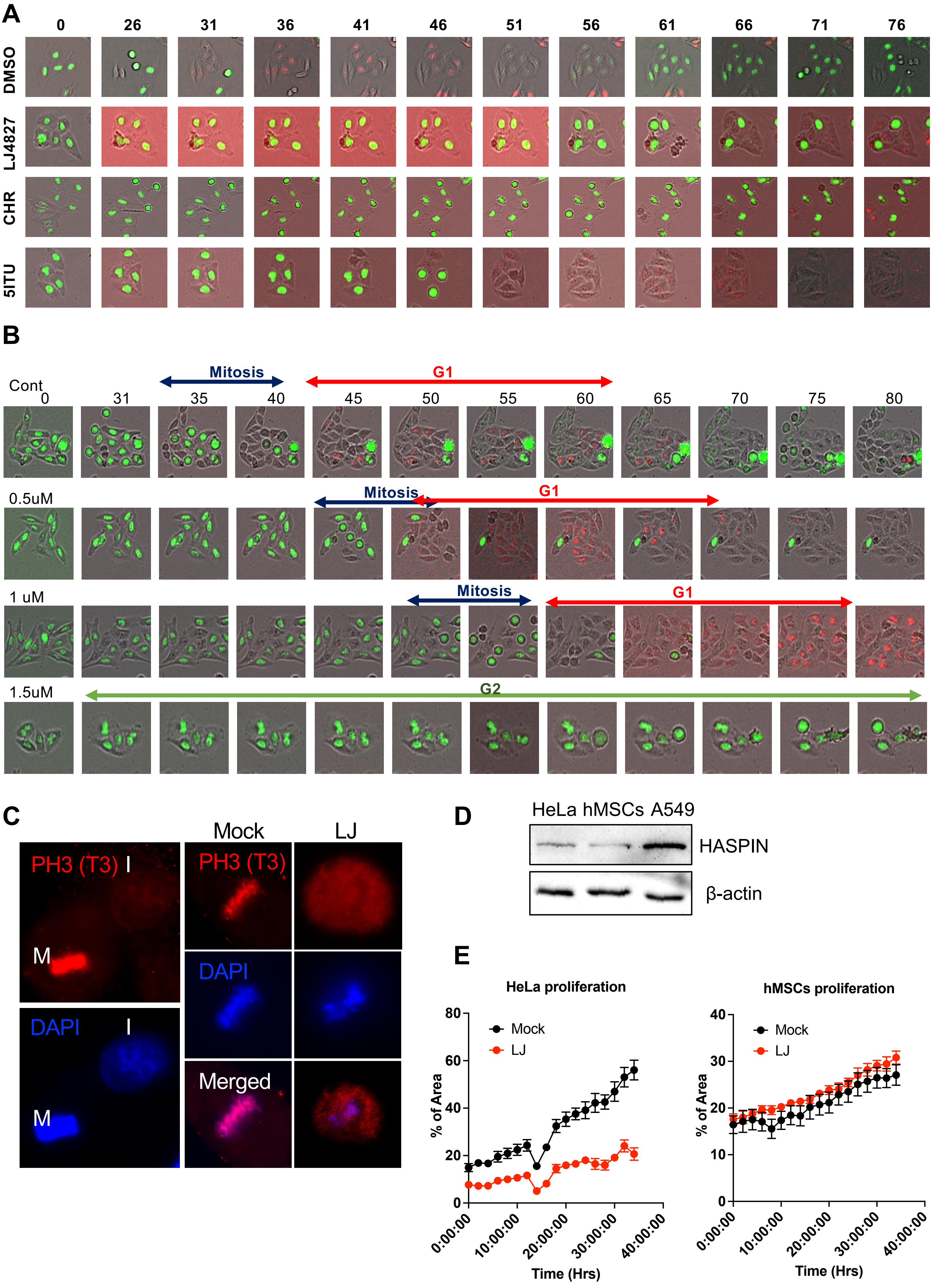

A
0 26 31 36 41 46 51 56 61 66 71 76
DMSO
LJ4827
CHR
5ITU
B
Cont
0 31 35 40 45 50 55 60 65 70 75 80
Mitosis
G1
0.5uM
Mitosis
G1
1 uM
Mitosis
G1
1.5uM
G2
C
PH3 (T3)
I
M
DAPI
I
M
Mock
LJ
PH3 (T3)
DAPI
Merged
D
HeLa hMSCs A549
HASPIN
β-actin
E
HeLa proliferation
Mock
LJ
% of Area
Time (Hrs)
0:00:00 10:00:00 20:00:00 30:00:00 40:00:00
hMSCs proliferation
Mock
LJ
% of Area
Time (Hrs)

**Figure S4**

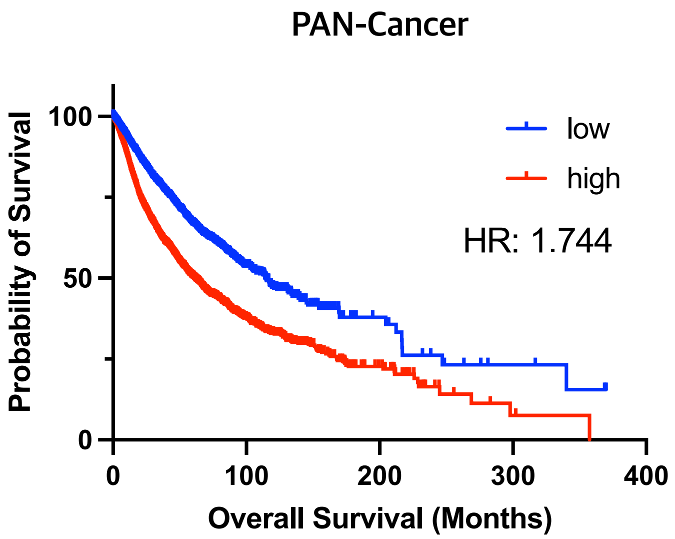

PAN-Cancer
Probability of Survival
100
50
0
low
high
HR: 1.744
0
100
200
300
400
Overall Survival (Months)

### Figure S5

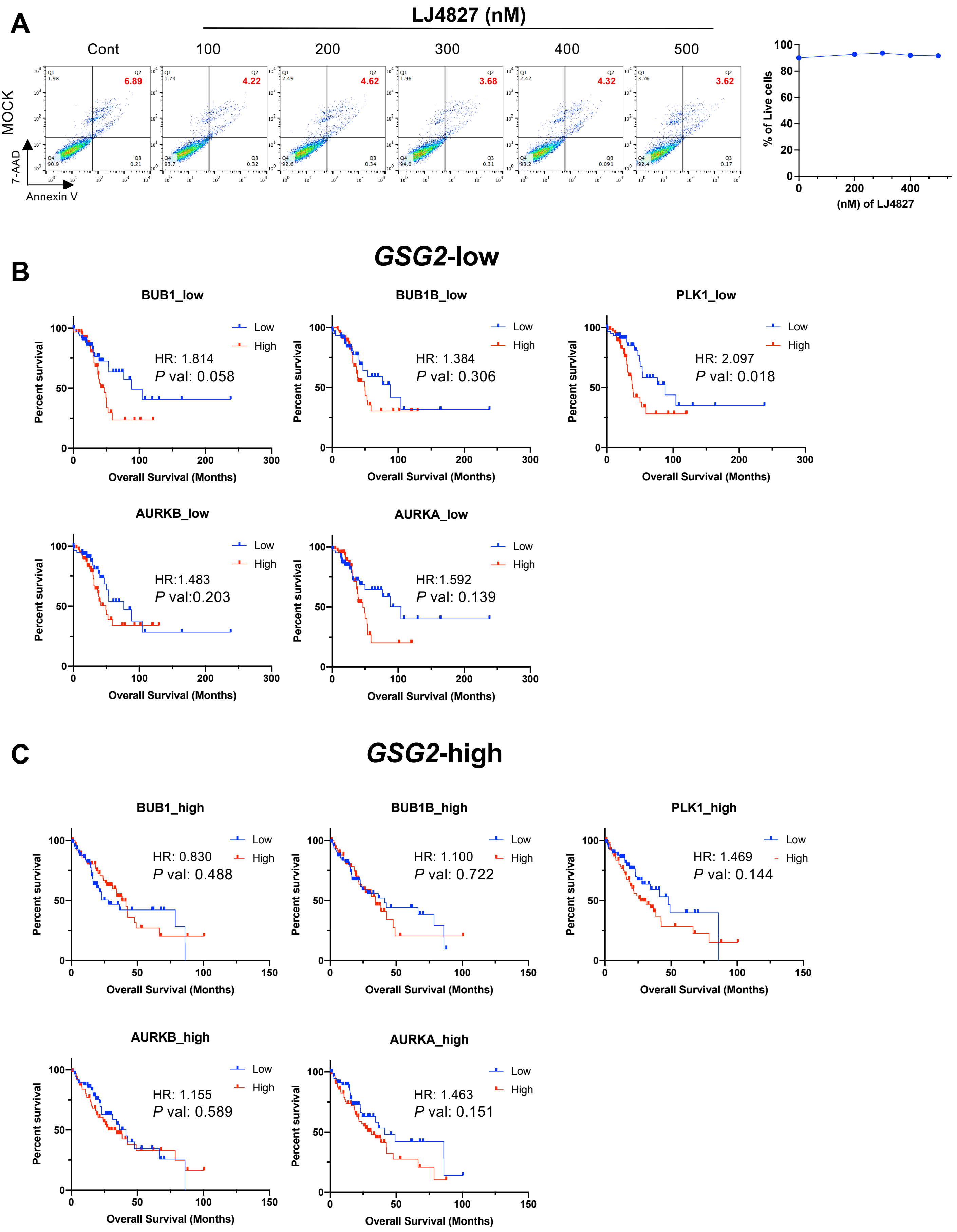

A
LJ4827 (nM)
Cont
100
200
300
400
500
MOCK
7-AAD
Annexin V
6.89
4.22
4.62
3.68
4.32
3.62
% of Live cells
(nM) of LJ4827
B
GSG2-low
BUB1_low
HR: 1.814
P val: 0.058
BUB1B_low
HR: 1.384
P val: 0.306
PLK1_low
HR: 2.097
P val: 0.018
AURKB_low
HR:1.483
P val:0.203
AURKA_low
HR:1.592
P val: 0.139
C
GSG2-high
BUB1_high
HR: 0.830
P val: 0.488
BUB1B_high
HR: 1.100
P val: 0.722
PLK1_high
HR: 1.469
P val: 0.144
AURKB_high
HR: 1.155
P val: 0.589
AURKA_high
HR: 1.463
P val: 0.151
Low
High
Percent survival
Overall Survival (Months)

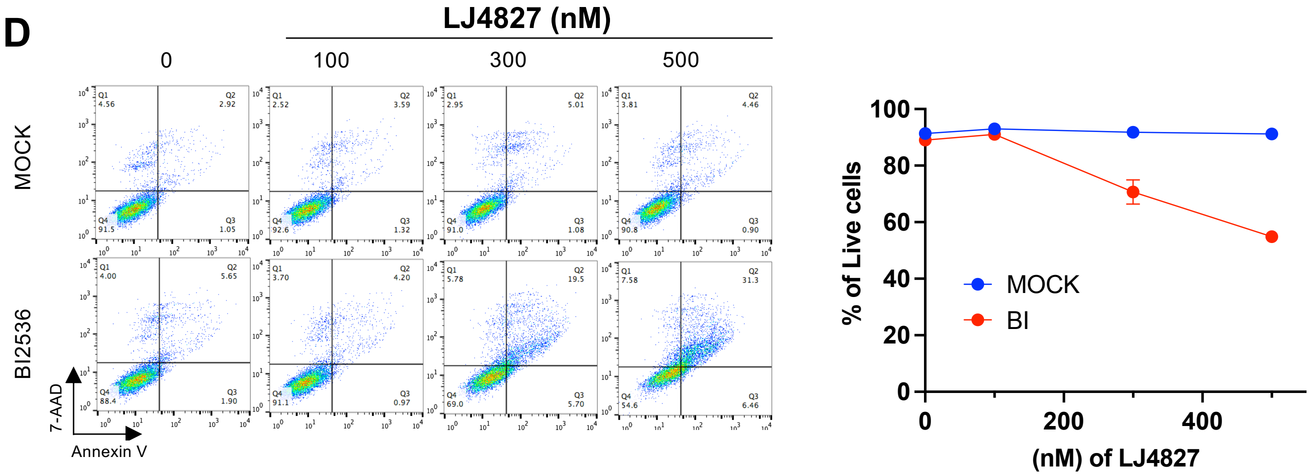
D
LJ4827 (nM)
0
100
300
500
MOCK
BI2536
7-AAD
Annexin V
Q1
Q2
Q3
Q4
4.56
2.92
91.5
1.05
2.52
3.59
92.6
1.32
2.95
5.01
91.0
1.08
3.81
4.46
90.8
0.90
4.00
5.65
88.4
1.90
3.70
4.20
91.1
0.97
5.78
19.5
69.0
5.70
7.58
31.3
54.6
6.46
% of Live cells
(nM) of LJ4827
MOCK
BI
